# A role for TRIP12 in regulating the E3 ligase network under proteotoxic stress

**DOI:** 10.64898/2026.11.10.764862

**Authors:** Hai Trinh, Farnusch Kaschani, Markus Kaiser, Smiths S. Lueong, Jens Siveke, Annette Paschen, Doris Hellerschmied, Michael Ehrmann

**Affiliations:** Center of Medical Biotechnology, Faculty of Biology, University of Duisburg-Essen, Universitaetsstr. 2, 45141 Essen, Germany; Bridge Institute of Experimental Tumor Therapy and DKTK Division of Solid Tumor Translational Oncology, West German Cancer Center, University Hospital Essen, University of Duisburg-Essen, Essen, Germany; German Cancer Consortium (DKTK), partner site Essen, a partnership between German Cancer Research Center (DKFZ) and University Hospital Essen, Germany; Department of Dermatology, University Hospital Essen, University of Duisburg-Essen, Essen, Germany

**Keywords:** HSP90, TRIP12, proteostasis, melanoma, E3 ligases, quantitative proteomics

## Abstract

Cellular proteostasis relies on the coordinated action of molecular chaperones and protein degradation pathways to maintain protein folding and turnover. While the roles of chaperones and the proteasome are well established, how ubiquitin E3 ligases contribute to the proteostasis network remains incompletely understood. Here, we analyzed proteostasis in melanoma cells using quantitative proteomics under mild protein folding stress induced by HSP90 and proteasome inhibition.

We identified the HECT-type E3 ligase TRIP12 as a proteostasis-responsive factor that was upregulated upon proteasome inhibition. Depletion of TRIP12 led to protein accumulation of a distinct group of epigenetic and transcriptional regulators, suggesting that these proteins may constitute a novel class of TRIP12 targets. Notably, the loss of TRIP12 led to a widespread reduction in the abundance of many other proteins, including components of the proteostasis network, particularly ubiquitin E3 ligases. These changes occurred largely without corresponding mRNA alterations, indicating post-transcriptional regulation.

HSP90 inhibition similarly reduced protein abundance and acted in an additive manner with TRIP12 depletion to further destabilize the proteome. Together, our findings suggest that TRIP12 functions as a regulator of the E3 ligase network and proteostasis capacity, and that its loss, particularly under folding stress, causes a reduction of global protein content. These results reveal an unexpected role for TRIP12 in maintaining proteome stability and support the functional interplay between ubiquitin E3 ligases and molecular chaperones in buffering proteotoxic stress.

## Introduction

Cellular protein quality control relies on protein repair and degradation that are performed by molecular chaperones and proteases, respectively^1–3^. Molecular chaperones assist the folding of nascent proteins, the refolding of misfolded proteins, and in delivering damaged proteins to proteases for degradation^4^. There are more than 300 genes encoding various classes of chaperones and co-chaperones in the human genome^5^. The most prominent chaperones are heat shock proteins, many of which are ATP-dependent folding factors^5^. HSP90 is one of the most important and abundant chaperones assisting in protein folding and refolding and has been shown to buffer primary mutations until the emergence of functional suppressor mutations during evolution^6^. HSP90 activity is essential for cancer cells, which carry a high burden of missense mutations and rely on many oncogenic drivers that are HSP90-sensitive client proteins; this dependency makes HSP90 a prominent target for cancer therapy^6,7^. On the other hand, the human genome encodes nearly 600 genes for proteases^8,9^. Proteases are classified according to the amino acid residue or ion that performs the chemical reaction i.e., the hydrolysis of the peptide bond. There are five major classes, the serine, metallo, cysteine, aspartic and threonine proteases, of which the threonine proteases have the fewest members. Curiously, although the proteasome is one of the most abundant and important proteases in the cell, it belongs to the threonine proteases^10^. The proteasome is an essential element of the ubiquitin-proteasome system, which is critical for targeted protein degradation in mammalian cells^11^. The process of targeted protein degradation is typically initiated by the covalent attachment of polyubiquitin chains to lysine residues of a target protein^12^, and the substrate specificity of this process is dictated by ubiquitin E3 ligases. The human genome encodes more than 600 E3 ligases, which are currently grouped into four major families: HECT (homologous to E6-AP C-terminus), RBR (RING-in between-RING), RCR (RING Cys relay), RING (really interesting new gene) ligases^13^. These classes differ in their interactions with E2 conjugating enzymes, their mechanisms of substrate binding and ubiquitin transfer, and the specific peptide motifs they recognize within their substrates^13^. While the proteasome is mechanistically well studied, much remains to be learned about the substrate specificity, regulation, and functional relevance of many E3 ligases^12^.

In this study, we aimed at identifying proteostasis factors that respond to protein folding stress by using two melanoma cell lines that harbor a large number of missense mutations, making them a suitable model for cells with a high basal folding burden and a strong reliance on protein folding and degradation pathways. These cell lines occupy opposite ends of the melanoma differentiation spectrum, allowing us to capture responses of more general relevance. To facilitate the identification of responsive protein quality control factors, we further perturbed the model system using low-dose inhibition of HSP90 and the proteasome, which are key components of the protein folding and degradation pathways. This approach allowed us to identify the most sensitive and responsive factors in protein quality control, which may not be apparent at higher concentrations of inhibitors. Proteomic analysis identified the E3 ligase TRIP12 as upregulated under protein folding stress caused by inhibition of the proteasome in one of the melanoma cell lines used in this study. TRIP12 has been implicated in diverse cellular processes, including DNA damage repair, chromatin regulation, cell differentiation, cell-cycle progression, and recently in oxidative stress^14,15^. To further investigate its role in protein folding stress, we performed proteomic studies in TRIP12-depleted cells. Our results suggest a new role of TRIP12 in regulating epigenetic regulators and transcriptional regulators, many of which may be rapidly turned over under normal conditions by TRIP12. Additionally, the data suggest that TRIP12 might play a role in regulating the proteostasis network, in particular E3 ligases, and may also act together with HSP90 to modulate the E3 ligase network.

## Results

### Defining proteotoxic stress conditions for quantitative proteomics experiments

To identify proteostasis factors responsive to proteotoxic stress, we used two melanoma cell lines that occupy opposite ends of the melanoma differentiation spectrum. MaMel102 is a differentiated melanoma cell line, expressing high levels of melanocyte-inducing transcription factor (MITF) and melanoma antigen recognized by T-cells 1 (MLANA), whereas MaMel94 is a de-differentiated melanoma cell line with significantly lower levels of MITF and MLANA (**Fig. S1A**)^16,17^.

Both cell lines carry a substantial load of missense mutations, with 678 and 1123 mutations in MaMel94 and MaMel102, respectively **(Fig. S1A)**^18^. As a result, these cell lines are expected to experience a high basal folding burden and to rely heavily on HSP90 and the proteasome for proteostasis. To further increase protein folding stress, we applied inhibitors of the HSP90 chaperone i.e., 17-Dimethylaminoethylamino-17-demethoxygeldanamycin (17-DMAG)^19^ and of the proteasome i.e., Bortezomib (BTZ)^20^.

To find conditions under which HSP90 and the proteasome are inhibited without compromising cell viability, we first assessed the effects of 17-DMAG and BTZ, using a 3-(4,5-Dimethylthiazol-2-yl)-5-(3-Carboxymethoxyphenyl)-2-(4-sulfophenyl)-2H-Tetrazolium (MTS) assay, which quantifies cellular metabolic activity as an indicator of cell viability. Treatment with 17-DMAG at concentrations between 10 nM and 1000 nM for 24 hours resulted in a modest reduction in cell viability, ranging from 5% to 20%, in both MaMel94 and MaMel102 cell lines (**Fig. 1A**, left panel). Similarly, extended treatment for 48 hours with 17-DMAG did not significantly impact cell viability at concentrations below 100 nM in either cell line (**Fig. 1A**, left panel). However, at concentrations of 200 nM and above, a more pronounced reduction in cell viability was observed. MaMel94 exhibited a steep decline in cell viability, with a 90% reduction at 1000 nM, whereas MaMel102 showed a more gradual decrease, resulting in a 40% reduction at 1000 nM (**Fig. 1A**, left panel). Western blot analysis of CDK6, an HSP90 client protein^21,22^ revealed a concentration-dependent decrease in protein levels after 24-hour treatment, with significant effects observed at concentrations of 250 nM and above (**Fig. S1B**), confirming the inhibition of HSP90. In contrast to 17-DMAG, BTZ treatment had a more pronounced effect on cell viability within 24 hours (**Fig. 1A**, right panel). The MTS assay revealed IC_50_ values of 26 nM for MaMel94 and 59 nM for MaMel102 after 24 hours of treatment with BTZ. Notably, proteasomal activity was rapidly and dose-dependently inhibited by BTZ in both cell lines, with a decrease of 60% at 5 nM and of 90% at 10 nM after 8 hours of treatment (**Fig. S1C**, left panel). Furthermore, accumulation of ubiquitinated proteins was observed in both cell lines, with a slight increase at 5 nM BTZ and a more pronounced increase at 10 nM BTZ and above (**Fig. S1C**, right panel).

**Fig. 1.**
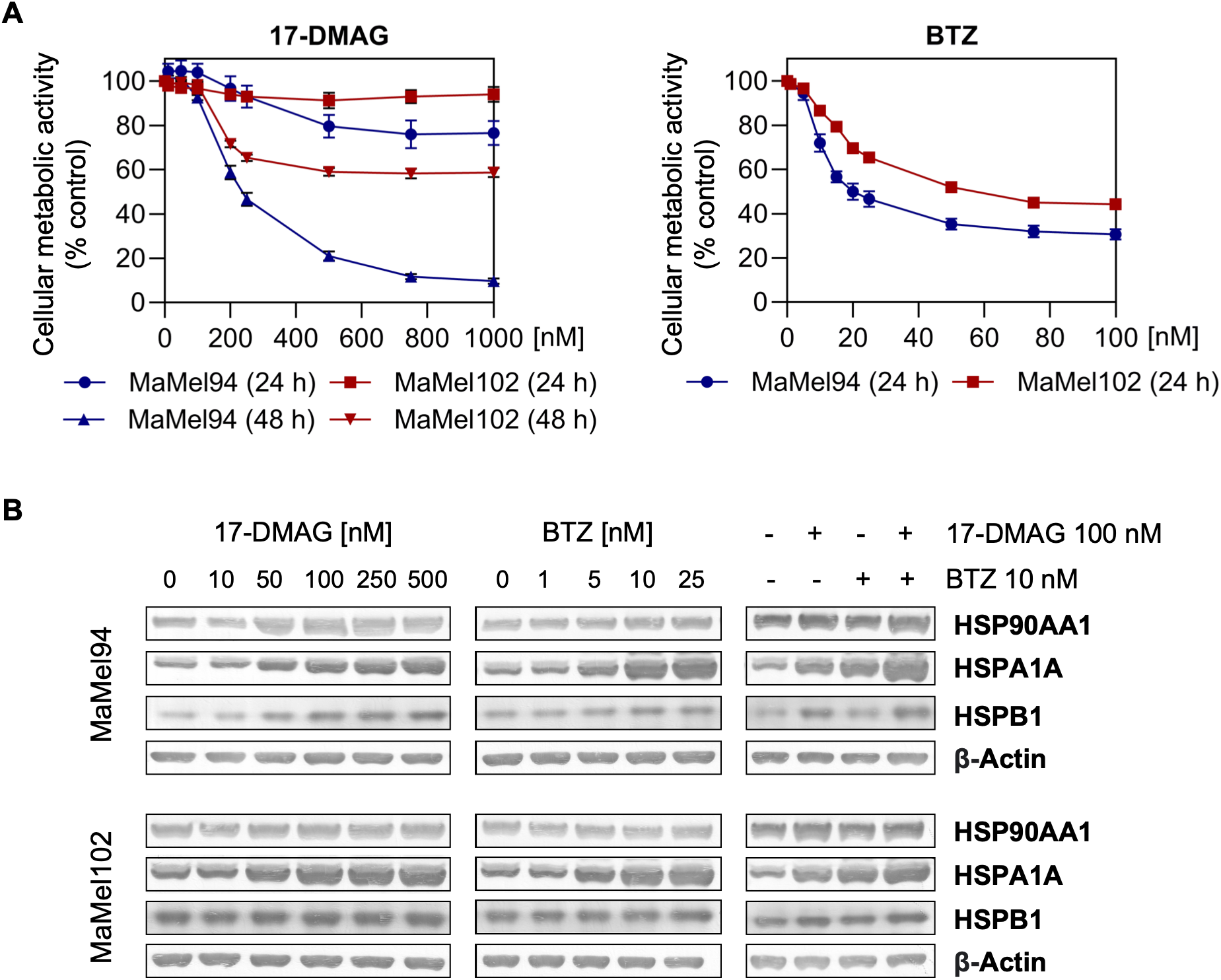
Induction of proteotoxic stress by the proteasome and HSP90 inhibitors. **(A)** Cell viability of MaMel94 and MaMel102 cells was assessed by MTS assay after treatment with Bortezomib (BTZ) for 24 h at concentrations ranging from 1 nM to 100 nM, or 17-DMAG for 24 h or 48 h at concentrations ranging from 10 nM to 1000 nM. Values are mean ± SD of three independent replicates (n = 3), normalized to DMSO control (=100%). **(B)** Western blot analysis of protein folding stress markers (HSP90AA1, HSPA1A and HSPB1) was performed on cells treated with 17-DMAG at concentrations ranging from 10 nM to 500 nM (left panel), BTZ at concentrations ranging from 1 nM to 25 nM (middle panel), or combination treatments 17-DMAG at 100 nM and/or BTZ at 10 nM (right panel), all for 24 h, with DMSO as vehicle control. β-Actin served as loading control for Western blot.

To confirm the induction of protein folding stress following HSP90 and the proteasome inhibition, we performed Western blotting to assess the protein levels of established heat shock markers, including HSP90AA1 (a stress-inducible form of HSP90), HSPA1A (a stress-inducible form of HSP70), and HSPB1 (HSP27). Our results show that both BTZ and 17-DMAG induced a concentration-dependent increase in HSPA1A level in both cell lines (**Fig. 1B**). Notably, the combination of inhibitors elicited a more pronounced upregulation of HSPA1A compared to single inhibitor treatment. Furthermore, HSPA1A upregulation was observed at remarkably low concentrations of BTZ (5 nM) and 17-DMAG (50 nM) (**Fig. 1B**), at which cellular metabolic activity remained largely intact (**Fig. 1A**). This suggests that protein folding stress can be triggered by subtle disruptions of proteostasis, even at dosages that do not significantly compromise cellular viability. These low concentrations therefore allow us to study protein folding stress in cells that remain relatively healthy.

### Proteomic profiles of cells treated with BTZ, 17-DMAG, or their combination

To investigate the changes of the proteomes of MaMel94 and MaMel102 cells in response to protein folding stress in an unbiased manner, we performed label-free quantitative proteomics experiments using low concentrations of proteasome and HSP90 inhibitors i.e., 5 nM BTZ and 50 nM 17-DMAG, alone or in combination (**Fig. 2A**). The analysis identified 3575 and 3924 proteins in MaMel94 and MaMel102 cells, respectively, with 3291 proteins identified in both cell lines (**Fig. S2A**, left panel). Furthermore, the proteomic profiles of the two cell lines reflected their distinct differentiation states, with melanoma differentiation markers (MITF, MLANA, PMEL, TYR, and SOX10) detected exclusively in MaMel102 samples, and markers of de-differentiated melanoma (AXL, EGFR, and TGFB1I1) present exclusively in MaMel94 samples (**Fig. S2A**, right panel)^16,17,23^. Proteins were defined as upregulated or downregulated based on a log2 fold change ≥ log2(1.3) or ≤ −log2(1.3), respectively, and an adjusted p-value < 0.05. In response to 17-DMAG treatment, 21 and 18 proteins were upregulated, and 4 and 20 proteins were downregulated in MaMel94 and MaMel102 cells, respectively (**Fig. 2B**). In response to BTZ treatment, 126 and 151 proteins were upregulated, and 112 and 87 proteins were downregulated in MaMel94 and MaMel102 cells, respectively (**Fig. 2B**). The combination of BTZ and 17-DMAG resulted in 51 upregulated proteins and 19 downregulated proteins in MaMel94, and 275 upregulated proteins and 163 downregulated proteins in MaMel102 cells (**Fig. 2B**). To identify biological processes that were affected by 17-DMAG and BTZ treatment, we performed gene ontology (GO) enrichment analysis of differentially abundant proteins in each treatment condition. The analysis revealed that biological processes related to proteostasis, including “response to heat”, “protein folding”, “regulation of protein stability” and “response to unfolded protein”, were significantly enriched in response to 17-DMAG treatment in both MaMel94 and MaMel102 cell lines (**Fig. S2B**). Similarly, the biological processes “response to heat”, “response to topologically incorrect protein” and “chaperone cofactor-dependent protein refolding” were also enriched following BTZ treatment (**Fig. S2B**). These results confirm that 17-DMAG and BTZ, even at low concentrations, induce protein misfolding stress in the two cell lines, triggering a cellular response aimed at mitigating the effects of protein misfolding and maintaining proteostasis.

**Fig. 2.**
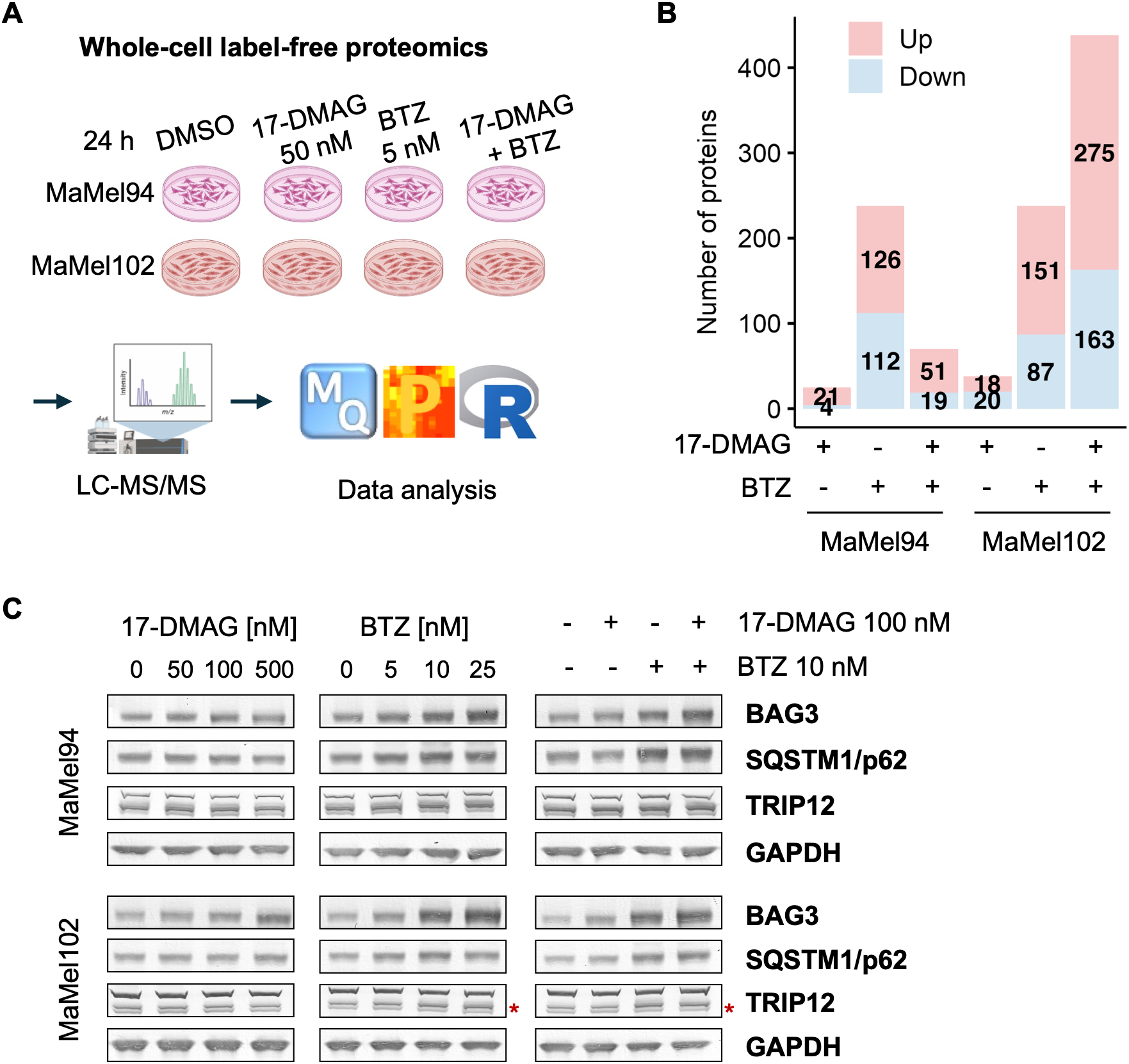
A low-dose inhibitor approach combined with whole-cell proteomics to investigate the effects of proteasome and HSP90 inhibition. **(A)** Schematic representation: cells were treated with 17-DMAG (50 nM) and/or BTZ (5 nM) for 24 h and analyzed by label-free quantification using LC-MS/MS (n = 4 biological replicates). **(B)** The numbers of upregulated (pink) and downregulated (blue) proteins in each treatment condition (compared with DMSO control) were identified from unpaired two-tailed t-tests, with a fold-change threshold of |log2FC| ≥ log2(1.3) and a permutation-based adjusted p-value < 0.05. **(C)** Validation of protein levels via Western blotting of selected proteins (BAG3, SQSTM1, TRIP12) after treatment of cells with various concentrations of 17-DMAG and/or BTZ for 24 h, using GAPDH as a loading control. Red asterisks indicate the lower TRIP12 band that is upregulated in BTZ and combination treatments in MaMel102.

In response to 17-DMAG, we identified 11 proteins that were upregulated in both MaMel94 and MaMel102 cell lines, all of which are members of various heat shock protein (HSP) families (**Fig. S3A**, left panel). These include HSP90 (HSP90AA1, HSP90AB1), HSP70 (HSPA1A, HSPA8), HSP27 (HSPB1), and HSP110 (HSPH1), as well as HSP70 co-chaperones (BAG3, DNAJA1, and DNAJB1) and HSP90 co-chaperones (AHSA1 and CHORDC1) (**Fig. S3A**, left panel). The consistent detection of these proteins across all replicates and samples in both cell lines suggests that they play crucial roles in cellular function. These findings indicate that cells attempt to mitigate the effects of HSP90 inhibition by upregulating HSP70 and its associated co-chaperones, together with other heat shock proteins, a phenomenon that aligns with previously reported observations^24,25^.

In response to BTZ, we identified 28 proteins that were upregulated in both MaMel94 and MaMel102 cell lines (**Fig. S3A**, middle panel). These upregulated proteins belong to multiple functional classes, including chaperones and co-chaperones of HSP70 (HSPA1A, BAG3, DNAJA1), protein ubiquitination pathways (SQSTM1/p62, UBXN4, E3 ligases), redox regulation (APOL2, BLVRB, GCLM, SRXN1, TXNDC9), vesicular-related proteins (CHMP5, CLTB, TRAPPC3), and mRNA-related proteins (SUPT6H, GRB2, DNTTIP2, PELO). The upregulation of these proteins suggests that, in response to the compromised proteasomal degradation capacity induced by BTZ, cells activate a multifaceted response to maintain protein homeostasis, involving increased chaperone production, enhanced autophagic degradation of misfolded proteins and induction of redox regulatory factors, which is consistent with existing literature^26–29^.

As E3 ligases play a crucial role in the ubiquitin-proteasome system, we analyzed the protein levels of E3 ligases in MaMel94 and MaMel102 cells in response to proteasome and HSP90 inhibition. Our analysis revealed 15 E3 ligases that were present at elevated levels in at least one treatment condition in at least one cell line (**Fig. S3B**). Notably, none of these E3 ligases were upregulated in response to 17-DMAG treatment alone, whereas 3 out of 55 E3 ligases detected in MaMel94 and 12 out of 55 E3 ligases detected in MaMel102 were upregulated in response to either BTZ or combination treatment (**Fig. S3B**, left panel). Among these, six E3 ligases were upregulated in more than one condition, including RING-type E3 ligases RMND5A, RNF181, UBR2, UBR3 and UBE4B, and HECT-type E3 ligase TRIP12 (**Fig. S3B**, left panel). Notably, TRIP12 and UBE4B were found to be abundant in both cell lines at basal levels (**Fig. S3B**, right panel). The abundance of TRIP12 and UBE4B in melanoma cells at basal levels and their upregulation in response to proteasome inhibition suggest a potential role in maintaining protein homeostasis under conditions of intrinsic protein folding stress. The recent implication of TRIP12 in the oxidative stress response^15^ further suggests that it may contribute to protecting cells against proteotoxic stress. Given that our melanoma cells serve as a model of intrinsic proteotoxic stress, TRIP12 emerges as a particularly interesting candidate to investigate further its potential role in maintaining protein homeostasis in this context. To validate the proteomics data, we performed Western blotting to assess the protein levels of BAG3, SQSTM1/p62, and TRIP12. Our proteomics analysis had shown that HSP70 co-chaperone BAG3 was upregulated in response to both 17-DMAG and BTZ treatments, whereas autophagy factor SQSTM1/p62 was upregulated in response to BTZ treatment but not to 17-DMAG treatment, and E3 ligase TRIP12 was upregulated in BTZ and combination treatment in MaMel102 (**Fig. S3C**). Western blot analysis confirmed these findings, showing a dose-dependent response of BAG3 to both treatments, with a stronger response to BTZ than 17-DMAG in both cell lines, and an additive effect in combination treatment; a dose-dependent increase of SQSTM1/p62 in response to BTZ, but no response to 17-DMAG; and upregulation of TRIP12 in MaMel102 cells in response to BTZ and combination treatment (**Fig. 2C**).

### Characterization of the HECT-type E3 ubiquitin ligase TRIP12

To investigate the role of TRIP12 in protein homeostasis, we performed siRNA-mediated knock-down in both melanoma cell lines. The TRIP12-targeting siRNA has been rigorously validated in several independent studies^30–33^ guaranteeing both specificity and potency. Western blot analysis confirmed efficient depletion of TRIP12 in both MaMel94 and MaMel102 cells (**Fig. 3A**). To further induce protein folding stress, we subsequently treated cells with 500 nM 17-DMAG. Western blot analysis confirmed that 17-DMAG treatment induced folding stress in both control and TRIP12-depleted cells, as evidenced by the robust upregulation of HSPA1A (**Fig. 3A**). To gain insights into the cellular response to TRIP12 depletion, HSP90 inhibition, and the combination of both conditions, we performed label-free quantitative proteomics analysis under these conditions.

**Fig. 3.**
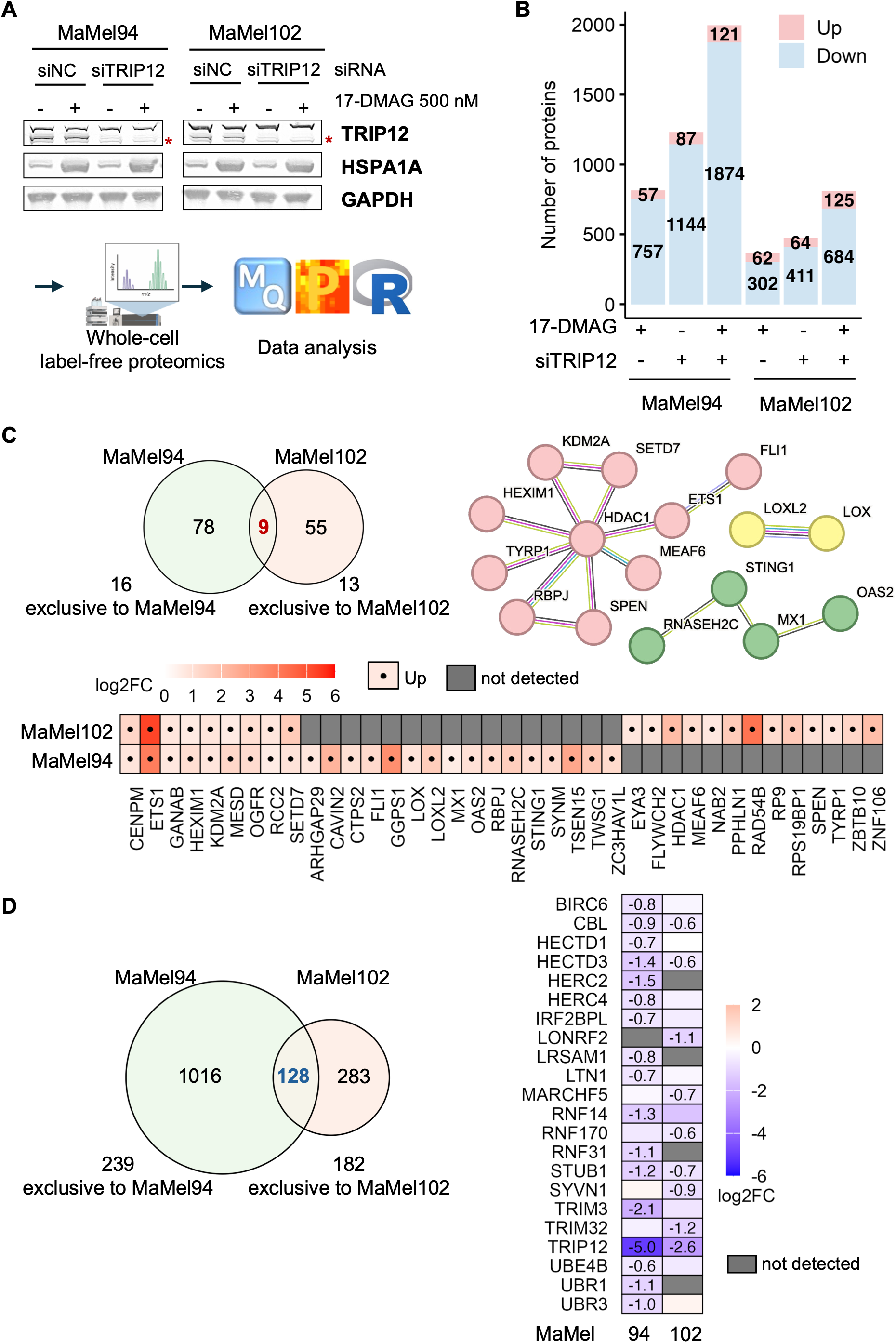
Proteomics analysis of the effects of TRIP12 depletion alone or in combination with HSP90 inhibition. **(A)** Cells were depleted of TRIP12 using siRNA for 72 h and subsequently treated with 17-DMAG (500 nM) for 24 h to induce protein folding stress. siNC and DMSO served as controls for siRNA transfection and 17-DMAG treatment, respectively. Western blot analysis confirmed TRIP12 depletion and induction of protein folding stress (HSPA1A), using GAPDH as a loading control. The same conditions were used for whole-cell label-free proteomics. **(B)** The numbers of upregulated (pink) and downregulated (blue) proteins in each treatment condition (compared with siNC + DMSO control) were identified using an unpaired two-tailed t-test with a fold-change threshold of |log2FC| ≥ log2(1.5) and a permutation-based adjusted p-value < 0.05. **(C)** Venn diagram shows the number of proteins upregulated in response to TRIP12 depletion in both cell lines, including overlapping proteins. Heatmap shows log2FC (TRIP12 depletion vs control) of proteins which were commonly upregulated in both cell lines, or upregulated in one cell line and absent in the other. Gray indicates protein not detected. STRING analysis of the proteins shown in the heatmap, with only connected proteins displayed. Three subnetworks are highlighted: epigenetic or transcriptional regulators (red), immune/viral response proteins (green), oxidases (yellow). **(D)** Venn diagram shows the number of proteins downregulated in response to TRIP12 depletion in both cell lines, including overlapping proteins. Heatmap shows log2FC of E3 ligases which were downregulated upon TRIP12 depletion in at least one cell line, with numerical values displayed only for downregulated protein (log2FC ≤-log2(1.5) and adjusted p-value < 0.05). Gray indicates protein not detected.

The proteomics analysis detected 4519 proteins in MaMel94, 4584 proteins in MaMel102, of which 4154 proteins were common to both cell lines. Proteins were defined as upregulated or downregulated based on a log2 fold change ≥ log2(1.5) or ≤ −log2(1.5), respectively, and an adjusted p-value < 0.05. In response to 500 nM 17-DMAG-mediated HSP90 inhibition, both cell lines exhibited a larger number of downregulated proteins than upregulated proteins, with 757 downregulated and 57 upregulated in MaMel94, and 302 downregulated and 62 upregulated in MaMel102 (**Fig. 3B**). Similarly, TRIP12 depletion resulted in 1144 proteins downregulated and 87 proteins upregulated in MaMel94, and 411 proteins downregulated and 64 proteins upregulated in MaMel102 (**Fig. 3B**). The combined perturbation of HSP90 inhibition and TRIP12 depletion had an additive effect, resulting in 1874 proteins downregulated and 121 proteins upregulated in MaMel94, and 684 proteins downregulated, and 125 proteins upregulated in MaMel102 (**Fig. 3B**). These results indicate that the combination of HSP90 inhibition and TRIP12 depletion leads to a more severe disruption of protein homeostasis in both cell lines.

### Downregulation of proteostasis factors following TRIP12 depletion

TRIP12 depletion resulted in the downregulation of 1144 proteins in MaMel94 and 411 proteins in MaMel102, with 128 proteins overlapping between the two cell lines (**Fig. 3D**). Gene Ontology biological process enrichment analysis of the downregulated proteins revealed that several of the top enriched terms are related to protein quality control in both cell lines (**Fig. S5A**). In MaMel94, the top 10 enriched pathways include “macroautophagy”, “protein folding” and “regulation of protein stability” (**Fig. S5A**). In MaMel102, the top 10 enriched pathways include “macroautophagy”, “ERAD pathway”, “Golgi vesicle transport”, and “response to endoplasmic reticulum stress” (**Fig. S5A**). Gene Ontology molecular function enrichment analysis further supported the impact of TRIP12 depletion on protein quality control. In MaMel94, the enriched terms include “ATP-dependent protein folding chaperone”, “proteasome binding”, “protein folding chaperone”, “ubiquitin-like protein ligase binding” and “unfolded protein binding” (**Fig. S5B**). Consistent with the gene ontology enrichment results, a closer inspection of the 128 commonly downregulated proteins in both cell lines upon TRIP12 depletion identified several key proteostasis factors, including the autophagy factor ATG7, ER-resident protein ERLEC1, folding factor FKBP1A, deubiquitinases (USP14, USP19, UFSP2), and E3 ligases (CBL, STUB1, HECTD3). Notably, the downregulation of these E3 ligases was unexpected, prompting a more detailed examination of the E3 ligase network upon TRIP12 depletion. Further analysis revealed a total of 16 E3 ligases in MaMel94 and 8 in MaMel102 to be downregulated upon TRIP12 depletion (**Fig. 3D**, right panel). Altogether, these data suggest that TRIP12 depletion might broadly perturb the protein quality control network, leading to a widespread downregulation of key proteostasis factors, including folding factors, autophagy factors, deubiquitinases, and notably, other E3 ligases.

To determine whether protein downregulation following TRIP12 depletion is driven by transcriptional changes, we performed bulk RNA sequencing and integrated the transcriptomic data with proteomic profiles. In MaMel94, only 32 proteins exhibited concurrent downregulation at both the protein and mRNA levels, whereas 1100 proteins were significantly reduced at the protein level without corresponding changes in transcript abundance (**Fig. S6B**). Similarly, in MaMel102, 95 proteins showed concordant downregulation at both levels, while 313 proteins were downregulated at the protein level without detectable changes in mRNA (**Fig. S6B**). These findings indicate that protein downregulation after TRIP12 depletion occurs mainly post-transcriptionally.

### Upregulation of epigenetic and transcriptional regulators following TRIP12 depletion

As TRIP12 is a HECT-type E3 ligase that primarily targets proteins for degradation^14^, we expected that its depletion would lead to the accumulation of its substrates and elevated levels of other E3 ligases that compensate for the loss of its activity. To test this model, we analyzed the proteins upregulated in response to TRIP12 depletion. In response to TRIP12 depletion, 87 proteins were upregulated in MaMel94, and 64 proteins were upregulated in MaMel102, with 9 proteins common to both cell lines (**Fig. 3C**). Notably, several of the upregulated proteins have been shown to either compensate for TRIP12 activity or be its targets. For example, the HECT-type E3 ligase UBR5, which increased in MaMel102 after TRIP12 depletion, has been reported to cooperate with TRIP12 in DNA-repair pathways^31,34^. Additionally, OTUD5, which accumulated in MaMel102 upon TRIP12 depletion, has been shown to be ubiquitylated by TRIP12 for proteasomal degradation^34^. Furthermore, STING, which was upregulated in MaMel94 after TRIP12 depletion, has been shown to be polyubiquitinated by TRIP12^35^. These observations confirm previously described TRIP12 targets and compensators, thereby supporting the validity of our dataset.

A closer examination of the 9 proteins upregulated in both MaMel94 and MaMel102 after TRIP12 depletion reveals their involvement in various cellular processes: chromatin regulation (SETD7, KDM2A), transcriptional regulation (HEXIM1, ETS1), chromosome segregation (CENPM, RCC2) and protein folding (GANAB, MESD) (**Fig. 3C**). In addition to these shared changes, we also observed cell-line-specific changes, with 16 proteins that were absent in MaMel102 but upregulated in MaMel94, and 13 proteins that were absent in MaMel94 but upregulated in MaMel102 upon TRIP12 depletion (**Fig. 3C**). These cell-line-specific proteins are involved in immune signaling (MX1 and STING1), RNA processing (RP9, TSEN15), DNA damage repair (RAD54B), chromatin regulation (HDAC1 and MEAF6), and transcriptional regulation (EYA3, FLI1, FLYWCH2, NAB2, RBPJ, SPEN, ZBTB10, ZNF106). A notable subset of these proteins displayed an “off-on” detection pattern: they were consistently undetected in control samples but detected across all replicates following TRIP12 depletion. This pattern was observed for CENPM and ETS1 in both cell lines, for GGPS1 and TSEN15 in MaMel94, and for HDAC1, MEAF6, PPHLN1, RAD54B and SPEN in MaMel102. Because imputation of log2-transformed LFQ intensities can obscure proteins consistently detected in one condition but not the other, we re-examined the raw, non-imputed LFQ intensity data and recovered additional proteins that were undetected in control samples but detected following TRIP12 depletion. The newly identified “off-on” proteins were predominantly epigenetic and transcriptional regulators. In MaMel94 cells, they included the transcription factor CUX1 and the histone methyltransferase NSD2. In MaMel102 cells, they included the transcriptional regulator DNTTIP1, the ETS-family transcription factor ELF1, the transcriptional coregulator ESS2, the epigenetic scaffold protein MEN1, and the zinc-finger transcription factor SP3. Examination of the transcriptomic profiles of genes encoding these “off-on” proteins showed that, with the sole exception of CENPM, which exhibited a modest increase in mRNA abundance in MaMel102 cells, transcript levels remained unchanged following TRIP12 depletion (Supplementary file, Sheet 4). The lack of transcriptional change led us to conclude that the accumulation of these “off-on” proteins after TRIP12 depletion is post-transcriptional; indicating that they are rapidly degraded via TRIP12-mediated activity under normal conditions. Some of these proteins may represent cell-line-specific substrates of TRIP12, and the prevalence of epigenetic and transcriptional regulators among them suggests that they might constitute a novel class of TRIP12 targets.

### Functional interaction between HSP90 and TRIP12

In response to the combined perturbation of HSP90 inhibition and TRIP12 depletion, a total of 121 proteins were upregulated, and 1874 proteins were downregulated in MaMel94, whereas 125 proteins were upregulated and 684 proteins were downregulated in MaMel102 (**Fig. 3B**). Notably, approximately half of the upregulated proteins in the combined treatment were not responsive to either single perturbation in both cell lines (**Fig. 4A**). Similarly, approximately one-third of the downregulated proteins in the combined perturbation were not responsive to either single perturbation (**Fig. 4A**). These data indicate that the combined perturbation imposes a greater stress, resulting in a markedly larger response compared to either single perturbation alone.

**Fig. 4.**
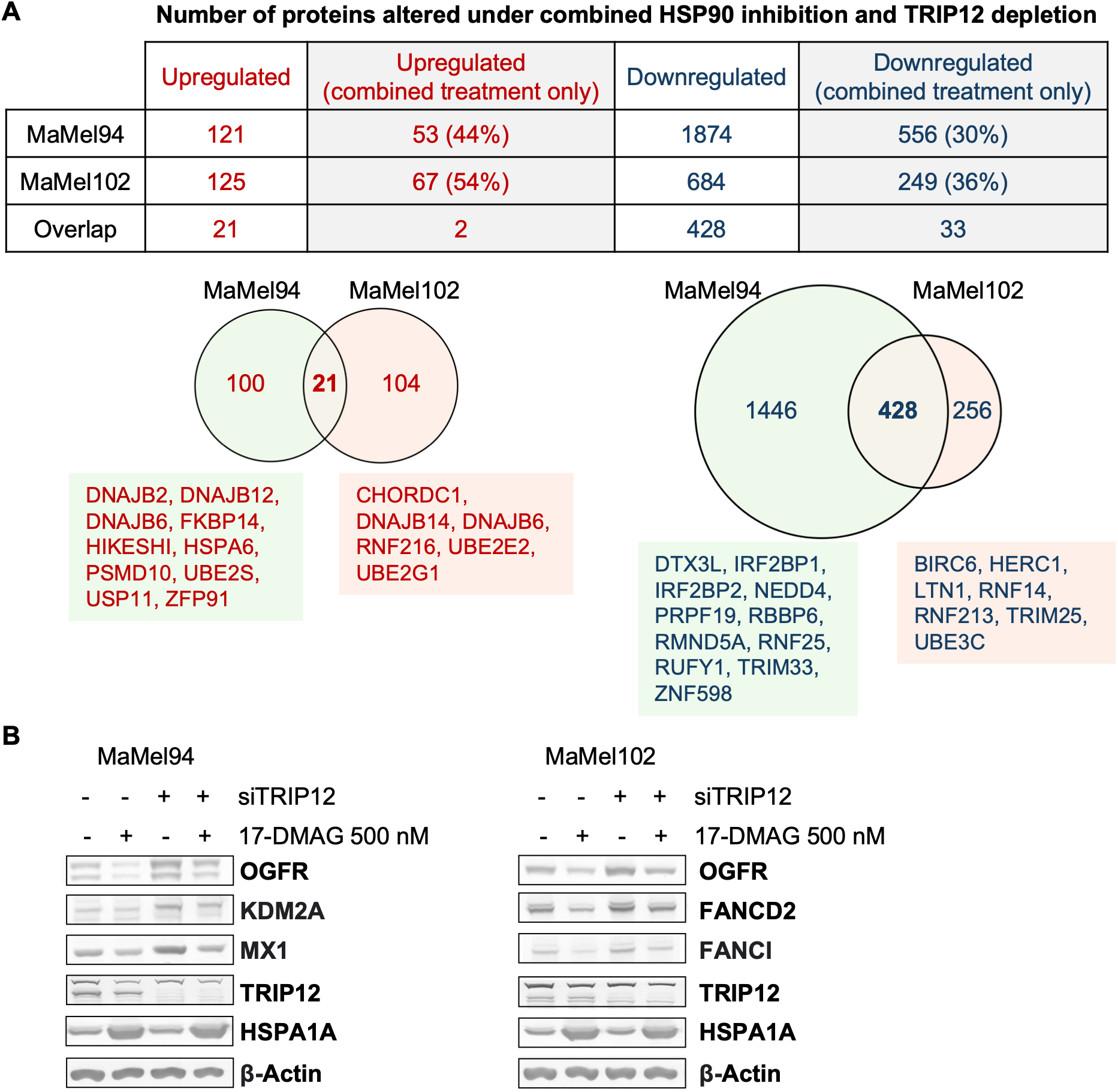
Combined effects of TRIP12 depletion and HSP90 inhibition. **(A)** Table and Venn diagrams show proteins upregulated and downregulated in response to combined HSP90 inhibition and TRIP12 depletion in MaMel94 and MaMel102. Proteins uniquely altered in the combined perturbation (but not in either single perturbation) are highlighted in the table, along with their percentage of the total number of upregulated or downregulated proteins. Several proteostasis factors are listed for each cell line below the Venn diagram. **(B)** Western blot validation of proteins potentially co-regulated by TRIP12 and HSP90. The proteins of interest were OGFR, KDM2A, and MX1 in MaMel94 cells, and OGFR, FANCD2, and FANCI in MaMel102 cells. Cells were depleted of TRIP12 for total 72 h and treated with 17-DMAG (500 nM) for 24 h. siNC and DMSO served as controls for siRNA transfection and 17-DMAG treatment, respectively. β-Actin was used as a loading control. TRIP12 and HSPA1A were assessed to confirm TRIP12 depletion and 17-DMAG-induced folding stress, respectively.

Notably, several proteins upregulated exclusively in the combined perturbation were components of the proteostasis network, including chaperones, co-chaperones, and ubiquitin-proteasome system members. In MaMel94, these proteins included DNAJB2, DNAJB12, DNAJB6, FKBP14, HIKESHI, HSPA6, PSMD10, UBE2S, USP11 and ZFP91 (**Fig. 4A**). Similarly, in MaMel102, they included 6 proteins, namely CHORDC1, DNAJB14, DNAJB6, RNF216, UBE2E2 and UBE2G1 (**Fig. 4A**).

Among the proteins downregulated exclusively in response to the combined perturbation, 11 E3 ligases were identified in MaMel94, i.e., DTX3L, IRF2BP1, IRF2BP2, NEDD4, PRPF19, RBBP6, RMND5A, RNF25, RUFY1, TRIM33, and ZNF598 (**Fig. 4A, Fig. S7A**). Similarly, 7 E3 ligases were downregulated exclusively in the combined perturbation in MaMel102, comprising BIRC6, HERC1, LTN1, RNF14, RNF213, TRIM25, and UBE3C (**Fig. 4A, Fig. S7A**). Interestingly, two of these E3 ligases (BIRC6 and LTN1) were also downregulated in all conditions in MaMel94, with a greater magnitude of downregulation in the combined perturbation compared to each single perturbation of HSP90 inhibition alone or TRIP12 depletion alone (**Fig. S7A**). Altogether, these findings reveal that simultaneous HSP90 inhibition and TRIP12 depletion elicits a broader and more pronounced effect on the E3 ligase network than either perturbation alone, suggesting a potential functional interplay between the chaperone HSP90 and the E3 ligase TRIP12 in maintaining E3 ligase homeostasis.

Another functional interplay of interest involves the proteins that are potentially co-regulated by both HSP90 and TRIP12. HSP90 stabilizes client proteins, and its inhibition can expose destabilized clients to ubiquitin-mediated proteasomal degradation involving E3 ligases like CHIP or CUL5^36–39^. We hypothesized that TRIP12 may function similarly, targeting HSP90 clients for degradation upon chaperone inhibition. To identify such proteins, we first selected those that were significantly downregulated upon HSP90 inhibition, indicating HSP90 client status. We then required that these proteins be rescued in the combined perturbation of HSP90 inhibition and TRIP12 depletion relative to HSP90 inhibition alone, suggesting that TRIP12 depletion prevents the degradation of destabilized clients. To further refine our search, we also looked for proteins that accumulated upon TRIP12 depletion under basal conditions, indicating that TRIP12 targets these substrates constitutively, not just under stress. This stringent filtering identified candidate proteins in both melanoma cell lines, including KDM2A, MX1, OAS2, OGFR, and RCC2 in MaMel94 (Supplementary file, Sheet 5), and FANCD2, FANCI, GINS3, NCAPD3, OGFR, PHIP, POLA1, RCC2 and UHRF1 in MaMel102 (Supplementary file, Sheet 5). To validate these proteomics findings, we performed Western blotting for several candidates: OGFR, KDM2A and MX1 in MaMel94, and OGFR, FANCD2, FANCI in MaMel102. Consistent with the proteomics data, protein levels decreased upon HSP90 inhibition compared to DMSO control in wild-type and also in TRIP12-depleted cells (**Fig. 4B**). Conversely, protein levels increased in TRIP12-depleted cells compared to controls, regardless of the presence or absence of HSP90 inhibition (**Fig. 4B**). These results indicate that TRIP12 depletion stabilizes these proteins, supporting a model in which TRIP12 contributes to the degradation of proteins normally stabilized by HSP90, particularly when chaperone function is compromised. These data suggest a functional interaction between HSP90 and TRIP12.

## Discussion

This study investigated how the proteomes of a differentiated and a de-differentiated melanoma cell line respond to disruptions in protein quality control caused by inhibition of the chaperone HSP90, inhibition of the 26S proteasome, and depletion of the E3 ligase TRIP12. While TRIP12 has been implicated in various cellular processes, including DNA damage repair, chromatin regulation, cell-cycle progression and recently in oxidative stress^14,15^, our data suggest that this E3 ligase may also function as an important regulator of proteostasis.

Analysis of our proteomic and transcriptomic datasets revealed that TRIP12 depletion led to widespread downregulation of proteins, with 1144 proteins downregulated in MaMel94 and 411 in MaMel102, predominantly at the post-transcriptional level (**Fig. 3B, Fig. S6B**). Gene Ontology biological process enrichment analysis of proteins downregulated following TRIP12 depletion identified several highly enriched terms related to protein quality control in both cell lines (**Fig. S5A**). Consistent with these findings, among the 128 proteins commonly downregulated across both cell lines were key proteostasis factors, including folding chaperones, autophagy-related proteins, deubiquitinases, and E3 ligases. Further analysis of E3 ligases revealed that, following TRIP12 depletion, 16 and 8 E3 ligases were downregulated in MaMel94 and MaMel102, respectively, whereas only one and three were upregulated in MaMel94 and MaMel102, respectively (**Fig. 3D, Fig. S7A**). This pronounced imbalance in E3 ligase abundance, coupled with the largely non-overlapping identities of affected E3 ligases between cell lines, suggests that TRIP12 does not merely target individual substrates for degradation, but might instead act as a regulator of the E3 ligase landscape itself.

We also examined the functional interplay between TRIP12 and HSP90 in maintaining proteostasis. Combined perturbation of HSP90 inhibition and TRIP12 depletion induced a greater proteotoxic burden than either intervention alone, as evidenced by a substantial increase in the number of differentially abundant proteins, many of which were uniquely altered under the combined perturbation (**Fig. 4A**). This additive effect suggests that HSP90 and TRIP12 may act in a coordinated manner to preserve protein homeostasis, and that simultaneous disruption of both systems overwhelms the cellular proteostasis capacity. A key point of functional convergence between TRIP12 and HSP90 appears to be the E3 ligase network. Our data showed that TRIP12 depletion led to the downregulation of multiple E3 ligases, indicating that TRIP12 plays a regulatory role in shaping the E3 ligase landscape (**Fig. 3D, Fig. S7A**). Independently, E3 ligases constitute a recognized class of client proteins for HSP90^22^. Consistent with this, our data showed that HSP90 inhibition reduced the abundance of 29 E3 ligases in MaMel94 and 6 in MaMel102 (**Fig. S7A**). Strikingly, the combined perturbation of TRIP12 depletion and HSP90 inhibition reduced the abundance of 11 E3 ligases in MaMel94 and seven in MaMel102 that were unaffected by either individual perturbation (**Fig. S7A**). Collectively, these findings reveal a coordinated impact on the E3 ligase network and open new avenues for investigating the coordinated regulation of E3 ligase activity by chaperones and E3 ligases themselves.

The unexpected observation that TRIP12 depletion resulted in reduced protein abundance, an effect further amplified by HSP90 inhibition and largely independent of transcriptional changes, may be explained by a TRIP12-dependent rewiring of the E3 ligase network. Loss of TRIP12 could permit compensatory ligases, such as UBR5 and UHRF1, to promote cross-and auto-ubiquitination events. Under conditions of HSP90 inhibition, increased conformational lability may further destabilize E3 ligases and promote their degradation. In parallel, stress-induced translational suppression, and destabilization of multi-protein complexes are also likely to contribute to a global shift toward protein loss. Together, these effects are more consistent with a collapse of proteostasis than with a simple substrate accumulation phenotype. Moreover, although the overall response to TRIP12 depletion and HSP90 inhibition was conserved across melanoma cell states, the specific E3 ligases affected were largely non-overlapping. Although an influence of cell line-specific mutations cannot be excluded, this observation suggests that the phenotype may arise from cell state-specific reconfiguration of the ubiquitin system rather than from TRIP12 acting directly on a defined set of substrates.

Our data further revealed that epigenetic and transcriptional regulators might constitute a novel class of potential TRIP12 substrates. Proteins including CENPM, CUX1, DNTTIP1, ELF1, ESS2, ETS1, GGPS1, HDAC1, MEAF6, MEN1, NSD2, PPHLN1, RAD54B, SP3, SPEN, and TSEN15 were consistently undetected in control samples but detected across all replicates following TRIP12 depletion (**Fig. 3C**). Notably, the mRNA levels of all these proteins, except CENPM, remained unchanged after TRIP12 loss (Supplementary file, Sheet 4), indicating that their accumulation occurs post-transcriptionally. This strongly suggests that these proteins are rapidly degraded under normal conditions and stabilized upon TRIP12 depletion, indicating that they may represent candidate substrates of TRIP12 and warrant further investigation. Furthermore, some of these proteins were detected only in one cell line, including GGPS1, NSD2 and TSEN15 in MaMel94, and HDAC1, MEAF6, PPHLN1, RAD54B, and SPEN in MaMel102 (**Fig. 3C**), suggesting that the substrate pool of TRIP12 may be cell-line-specific.

Whole-cell proteomics further revealed that even subtle perturbations of HSP90 and the proteasome were sufficient to trigger coordinated cellular stress responses. HSP90 inhibition with 17-DMAG at 50 nM induced a modest but specific response characterized by upregulation of heat shock proteins, including HSPA1A, HSPB1, and HSP90AA1, consistent with the established role of HSF1 in regulating heat shock gene expression following HSP90 inhibition (**Fig. S3A**, left panel)^40^. Upregulation of HSP90 co-activators such as AHSA1 and CHORDC1 likely reflects a compensatory attempt to restore HSP90 activity, while the concurrent induction of HSP70 and its co-chaperones, including DNAJA1 and DNAJB1, is consistent with the established cooperation between HSP70 and HSP90 in protein folding^24^. In contrast, proteasome inhibition with bortezomib elicited a more pronounced and complex response, including upregulation of the HSP70-HSP40-BAG3 complex, the autophagy receptor SQSTM1, and ERAD components UBXN4 and UFD1, reflecting activation of alternative protein degradation pathways to compensate for impaired proteasomal function (**Fig. S3A**, middle panel)^26,29,41^. Altogether, these findings validate the sensitivity of our low-dose inhibitor approach combined with quantitative proteomics for identifying the most responsive components and key regulatory factors of the proteostasis network in response to proteotoxic stress.

Beyond these shared responses to proteotoxic stress, the two melanoma cell lines also displayed distinct adaptive strategies. Notably, MaMel102 upregulated a larger set of proteasomal subunits and E3 ligases in response to bortezomib treatment than MaMel94 (**Fig. S3B, Fig. S8A**). This difference likely reflects distinct basal proteostasis states in the two cell lines. MaMel102 exhibited lower basal levels of proteasomal subunits compared with MaMel94, including components of the 19S regulatory particle (PSMC1/3/6 and PSMD2/5/10), the 20S core particle (PSMA1/2/3/6 and PSMB7/8), and the proteasome activator complex (PSME1 and PSME2) (**Fig. S8B**). In contrast, MaMel102 showed elevated basal levels of small heat shock proteins, such as HSPB1 and CRYAB, as well as the mitochondrial chaperonin HSPD1, compared with MaMel94 (**Fig. S8C**). Furthermore, the co-chaperones DNAJB12, DNAJB14 and DNAJC11 were more abundant in MaMel102, whereas DNAJA1, DNAJA2 and DNAJB11 were more abundant in MaMel94 (**Fig. S8C**). These findings are consistent with a recent study showing that melanoma tumors can employ diverse proteostasis strategies to cope with proteotoxic stress^42^. More broadly, such heterogeneity in proteostasis programs may have important therapeutic implications, as distinct tumor cell states may exhibit unique vulnerabilities to proteostasis-targeting drugs.

In conclusion, this study provides new insights into how melanoma cells respond to protein folding stress and identifies previously unrecognized functions of TRIP12. Our findings suggest that TRIP12 and HSP90 may act together to regulate the E3 ligase landscape and maintain proteostasis network balance. We further identify epigenetic and transcriptional regulators as a potential novel class of TRIP12 substrates, many of which are detected only upon TRIP12 depletion and appear to be cell-line-specific, suggesting context-dependent regulation. Future studies should aim to elucidate the mechanisms underlying the reduced protein abundance observed upon TRIP12 loss. The role of TRIP12 in potentially regulating epigenetic and transcriptional networks warrants further investigation into the molecular mechanisms underlying this function. Additional work is needed to define the molecular basis of the functional interaction between TRIP12 and HSP90, particularly in the coordinated regulation of E3 ligases and the co-regulation of potential shared substrates. Ultimately, targeting the TRIP12–HSP90 axis may offer a novel therapeutic strategy for disrupting proteostasis in melanoma cells.

## Materials and Methods

### Cell culture and treatments

Human melanoma cell lines (MaMel94 and MaMel102) were provided by Westdeutsche Biobank Essen (WBE/SCABIO, University Hospital Essen, University of Duisburg-Essen, Essen, Germany; approval no. SCABIO_114715). Cells were cultured in RPMI-1640 (Thermo Fisher Scientific, 21875091) supplemented with 10% v/v heat-inactivated Fetal Bovine Serum (Thermo Fisher Scientific, A5256801) and 1% v/v Penicillin/Streptomycin (Thermo Fisher Scientific, 15140122) at 37°C and 5% CO_2_. All the cells have tested negative for mycoplasma contamination. Cells were treated with 17-dimethylaminoethylamino-17-demethoxygeldanamycin (17-DMAG, Selleck Chemicals, S1142) and Bortezomib (BTZ, Sigma-Aldrich, 5043140001) at the concentrations indicated in the figure legends.

### siRNA transfection

For human TRIP12 knockdown, the following siRNAs (synthesized by Microsynth) were used: siTRIP12_s17810: 5’-GCAAUUUGAUUCGUUCAGA-3’; siNC_s229174: 5’-UUCUCCGAACGUGUCACGU-3’^31^. The siRNAs (final concentration 25 nM) were transfected into melanoma cells using Lipofectamine RNAiMAX Transfection Reagent (Thermo Fisher Scientific, 13778150) according to manufacturer’s instructions 72 h or 96 h prior to analysis. Subsequent treatment with 17-DMAG was performed 24 h before cell harvests. siRNA depletion efficiency was verified by Western blot.

### Cell viability assay (MTS Assay)

Cell viability was assessed using the CellTiter 96® AQueous One Solution Cell Proliferation Assay (Promega, G3581) according to the manufacturer’s protocol. Cells were seeded in 96-well plates (Greiner-Bio-One) and treated with different inhibitors as indicated in the figure legends. The absorbance was measured at 490 nm using a SpectraMax iD5 microplate reader (Molecular Devices) and the values were normalized to the DMSO-only control. All experiments were performed in biological triplicates.

### Proteasome activity assay

Cells were seeded in 384-well plates (Greiner Bio-One, 781098) in biological triplicates. After 48 h, when cells reached 90% confluency, they were treated with Bortezomib at the indicated concentrations or a DMSO-only solvent control for 8 h. At this time point, no changes in cell morphology or cell death were observed at any of the tested concentrations. Proteasome activity was measured using the Proteasome-Glo Chymotrypsin-Like Cell-Based Assay (Promega, G8660), which utilizes Suc-LLVY-aminoluciferin as a luminogenic substrate. Luminescence was then measured with a SpectraMax iD5 microplate reader (Molecular Devices). Data were normalized to the DMSO-only control group.

### Western blotting

Cells were lysed in ice-cold RIPA buffer containing 50 mM Tris-HCl (pH 7.4), 150 mM NaCl, 1% NP-40, 0.5% sodium deoxycholate, and 0.1% SDS, supplemented with protease cOmplete Protease Inhibitor Cocktail and PhosSTOP Phosphatase Inhibitor Cocktail (Roche). Lysates were centrifuged at 15000 × g for 30 min, and the protein concentration of the supernatant was determined using Pierce BCA protein assay kit (Thermo Fisher Scientific, 23227). Protein samples were separated by SDS-PAGE using 4–12% NuPAGE Bis-Tris precast gels (Thermo Fisher Scientific, WG1403BOX) and transferred onto PVDF membranes. The membranes were blocked with 3% BSA in TBST at room temperature for 1 h and incubated overnight at 4°C with primary antibodies diluted in blocking solution. The membranes were then incubated with alkaline phosphatase (AP)-conjugated secondary antibodies at room temperature for 2 h. Secondary antibodies used in this study include goat-α-rabbit IgG-AP conjugate (Thermo Fisher Scientific, A16113) and goat-α-mouse IgG-AP conjugate (Sigma-Aldrich, A1418). After incubation with secondary antibodies, signals were detected by colorimetric detection using the AP Substrate solution containing 100 mM Tris-HCl, pH 9.5, 100 mM NaCl, 5 mM MgCl_2_, 165 µg/ml BCIP, and 330 µg/ml NBT. The blots were scanned using CanoScan LiDE 400 (Canon) and analyzed using Phoretix 1D image analysis software. Details of primary antibodies are listed in Table S1.

### RNA extraction and qRT-PCR

Total RNA was extracted from MaMel94 and MaMel102 cells using the RNeasy Plus Mini Kit (Qiagen, 74134) according to the manufacturer’s instructions. DNase treatment was done using RNase-Free DNase Set (Qiagen, 79254). RNA concentration and purity were assessed using DS-11 Series Spectrophotometer (DeNovix). Complementary DNA (cDNA) was synthesized from 2 μg RNA using Verso cDNA Synthesis Kit (Thermo Fisher Scientific, AB1453B) and Anchored Oligo dT. qRT-PCR was performed using ABsolute QPCR SYBR Green Mix (Thermo Fisher Scientific, AB1158B) on a Bio-Rad CFX96 Touch Real-Time PCR Detection System. All primers were used at a final concentration of 400 nM. Gene expression levels were calculated using the ΔΔCt method (PMID: 18546601) and normalized to the housekeeping gene GAPDH and to treatment control. Primer sequences are listed in Table S2.

### RNA sequencing and data analysis

For RNA-Seq, total RNA was isolated from cells using the Maxwell® RSC simplyRNA Cells Kit (Promega, AS1390) according to manufacturer’s protocol, followed by polyA-enrichment using NEBNext Poly(A) mRNA Magnetic Isolation Module (New England Biolabs, E7490L). RNA sequencing libraries were generated using NEBNext® Ultra™ II Directional RNA Library Prep Kit Illumina (New England Biolabs, E7760S) and prepared and processed in a single batch with NEBNext® Multiplex Oligos for Illumina® (New England Biolabs, E7335L). Sequencing was performed on an Illumina NovaSeq 6000 platform using 100 bp paired-end reads, targeting a minimum depth of 60 million reads per sample. RNA seq experiments were conducted with four biological replicates. Paired-end FASTQ files were trimmed for adapter sequences and low-quality sequences using the Trim Galore tool, which integrates Cutadapt for adapter trimming and FastQC for quality control. The sequencing reads were then mapped to the human genome assembly GRCh38 and read counts per gene were quantified using the STAR aligner (PMID: 23104886). Only protein-coding genes were retained for downstream analysis. Lowly expressed genes were filtered out with edgeR v4.2.2 package (PMID: 39844453) using filterByExpr function with the experimental groups and a minimum count of ten (default setting). Differential expression analysis was performed with edgeR v4.2.2 package using glmQLFTest function. A gene was considered differentially expressed if the adjusted p-value (FDR) was < 0.05 and the absolute log2FC was ≥ log2(1.5).

### Sample preparation and clean-up for LC-MS/MS

For whole proteomes analysis, cells were harvested by trypsinization followed by twice wash with DPBS (Thermo Fisher Scientific, 14190144). Samples for proteomic analysis were prepared using the single-pot, solid-phase-enhanced sample-preparation (SP3) strategy^43^. All buffers and solutions were prepared with mass spectrometry-grade water (Avantor). Cell pellets were lysed in 50 mM HEPES (pH 8.0), 1% (w/v) SDS, 1% (v/v) NP-40, 10 mM TCEP, 40 mM chloroacetamide, and the samples were heated for 5 min at 90°C, then sonicated using Bioruptor (Diagenode) at high intensity 1 min ON/ 30 sec OFF for 10 min. Lysates were centrifuged at 20000 x g at room temperature for 20 min, and protein concentration of the supernatant was determined using Pierce 660 nm Protein Assay Reagent (Thermo Fisher Scientific, 22660) with the Ionic Detergent Compatibility Reagent (Thermo Fisher Scientific, 22663) according to the manufacturer’ instructions. Samples were stored at-20°C. Whole cell proteomes were prepared by treating 15 µg of total protein with 7 U of Benzonase (Merck Millipore, 71206) in dilution buffer (20 mM HEPES pH 8.0, 2 mM MgCl_2_) at 37°C, 1500 rpm for 30 min. Samples were then heated to 90°C for 5 min, cooled, and alkylated with 10 mM iodoacetamide at 37°C for 15 min. Proteins were then bound to a 1:1 mixture of hydrophilic and hydrophobic carboxylate modified Sera-Mag™ SpeedBeads (Cytiva, 45152105050250 and 65152105050250) using of an equal volume of pure ethanol (24°C, 10 min, 1000 rpm). The beads were washed with 80% ethanol and digested with a trypsin/LysC mix containing 0.6 μg of trypsin (Promega, V5111) and 0.6 µg LysC (FUJIFILM Wako Pure Chemical, 125-05061) in 25 mM ammonium bicarbonate at 37°C for 19 h while shaking at 1300 rpm. After digestion, clear solution containing the tryptic peptides were collected and then combined with the solution from an additional wash of the beads in 25 mM ammonium bicarbonate. This was followed by adding formic acid at 2% final concentration to inactivate trypsin.

Peptides were desalted on home-made C18 StageTips^44^ containing two layers of an octadecyl silica membrane (CDS Analytical). The StageTips were first activated and equilibrated by passing a series of solvents, including methanol, 80% (v/v) acetonitrile with 0.5% (v/v) formic acid. The acidified tryptic digests were then passed over the tips, followed by washing with formic acid to remove impurities. Bound peptides were eluted from the StageTips by two rounds of 25 μl 80% (v/v) acetonitrile with 0.5% (v/v) formic acid. The eluted peptides were then dried using a vacuum concentrator (Eppendorf) and the peptides were dissolved in 15 μl 0.1% (v/v) formic acid prior to analysis by mass spectrometry.

### LC-MS/MS settings

LC-MS/MS analysis of peptide samples was performed on an Orbitrap Fusion Lumos mass spectrometer (Thermo Fisher Scientific) coupled to an Easy nLC 1200 or to a Vanquish Neo ultra high-performance liquid chromatography (UHPLC) system (Thermo Fisher Scientific) that were operated in the one-column mode. The analytical column was a fused silica capillary (inner diameter 75 μm, outer diameter 360 µm, length 28 cm; New Objective) with an integrated sintered frit packed in-house with Kinetex 1.7 μm XB-C18 core shell material (Phenomenex). The analytical column was encased by a PRSO-V2 column oven (Sonation) and attached to a nanospray flex ion source (Thermo Fisher Scientific). The column oven temperature was set to 50°C during sample loading and data acquisition. The LCs were equipped with two mobile phases: solvent A (2% acetonitrile and 0.2% formic acid, in water) and solvent B (80% acetonitrile and 0.2% formic acid, in water). All solvents were of UHPLC grade (Honeywell). Peptides were directly loaded onto the analytical column with a maximum flow rate that would not exceed the set pressure limit of 950 bar and separated on the analytical column by running a 105 min gradient of solvent A and solvent B at a flow rate of 300 nl/min (start with 3% (v/v) B, gradient 3% to 6% (v/v) B for 5 min, gradient 6% to 29% (v/v) B for 70 min, gradient 29% to 42% (v/v) B for 15 min, gradient 42% to 100% (v/v) B for 5 min and 100% (v/v) B for 10 min). The mass spectrometer was controlled by the Orbitrap Fusion Lumos Tune Application and operated using the Xcalibur software.

### Peptide and protein identification using MaxQuant

Raw MS/MS spectra data were searched against the Uniprot H. sapiens reference proteome UP000005640_9606_OPPG (one protein per gene; 20606 entries) or UP000005640_9606_full (81803 entries) using Andromeda search in MaxQuant v2.5.2.0 with default settings^45,46^. Label-free quantification and match-between-runs were activated. All searches included a contaminants database search (as implemented in MaxQuant, 245 entries) and allowed for oxidation of methionine residues, acetylation of the protein N-terminus, and carbamidomethylation on Cysteine. Enzyme specificity was set to “Trypsin/P”, and the instrument type was set to Orbitrap. The precursor mass tolerance was set to ±20 ppm (first search) and ±4.5 ppm (main search), and the MS/MS match tolerance was set to ±0.5 Da. The peptide spectrum match FDR and the protein FDR were set to 0.01 (based on target-decoy approach). For protein quantification unique and razor peptides were allowed, and modified peptides were considered with a minimum score of 40. Label-free protein quantification was switched on, and unique and razor peptides were considered for quantification with a minimum ratio count of 2. Retention times were recalibrated based on the built-in nonlinear time-rescaling algorithm. MS/MS identifications were transferred between LC-MS/MS runs with the “match between runs” option in which the maximal match time window was set to 0.7 min and the alignment time window set to 20 min. The quantification is based on the “value at maximum” of the extracted ion current. At least two quantitation events were required for a quantifiable protein. Further analysis and filtering of the results was done in Perseus (v1.6.10.0)^47^.

### Proteomics data analysis with Perseus

The proteinGroup.txt files were imported to Perseus v1.6.10.0 for differential protein abundance analysis^47^. Protein groups were filtered to eliminate contaminants, reverse hits, and proteins identified by site only. Protein groups were retained only if they were quantified in at least 75% of the replicates for at least one sample condition across the two cell lines. Pearson-correlation heatmaps and PCA plots were generated to assess replicate reproducibility. The second replicate of MaMel102 in 17-DMAG treatment was excluded from subsequent analyses due to low correlation with other replicates. Before imputing missing values in each proteomics experiment, the data set was split into the MaMel94 and MaMel102 subsets, and each subset was filtered to retain only those protein groups that were quantified in at least 75% of the replicates for at least one sample condition in the respective cell line. Missing values were imputed with random values drawn from a normal distribution centered around the detection limit of the MS instrument (Perseus imputation; width = 0.3, down shift = 1.8).

Differentially abundant proteins (DAPs) were identified by two-tailed unpaired Student’s t-tests or Welch’s t-test (as appropriate) with a permutation-based false-discovery rate (FDR) of 0.05 and s0= 0 (default). Proteins were considered DAPs (upregulated or downregulated) when the adjusted p-value was < 0.05 and the absolute log2FC met the study-specific thresholds: ≥ log2(1.3) for the 17-DMAG and BTZ experiment, and ≥ log2(1.5) for the TRIP12 depletion and HSP90 inhibition experiment.

### Omics data analysis and visualization using R

Gene Ontology (GO) enrichment analysis was performed with the clusterProfiler R package (v4.12.6), using Gene Ontology Biological Process (GOBP) and Molecular Function (GOMF) gene sets from the org.Hs.eg.db R package (v3.19.1). Only the protein lists of interest were used as input for the enrichGO() function. Enrichment significance was assessed using the Benjamini–Hochberg method for multiple testing correction (adjusted p < 0.05, q-value < 0.02), and only gene sets containing between 10 and 500 genes were considered. To reduce redundancy among enriched GO terms, the simplify() function from clusterProfiler was applied (cutoff = 0.5, by = “p.adjust”). Venn diagrams were generated and plotted using the eulerr package (v7.0.2). All other plots for omics analyses were generated in R using the ggplot2 package (v3.5.1).

Reference gene lists used in this study are as follows: Somatic missense mutations in the MaMel94 and MaMel102 melanoma cell lines were identified from whole-exome sequencing data^18^, filtered by tumor barcode. Interactors of HSP90 isoforms (HSP90AA1, HSP90AB1, HSP90B1, TRAP1) were retrieved from the PicardLab HSP90 Chaperone Machine database (https://www.picard.ch/Hsp90Int/), restricted to Homo sapiens (taxon ID: 9606) and Interaction type of experimental. Proteins annotated as part of the cellular proteostasis network were obtained from Proteostasis Network Annotation 2.0 (https://www.proteostasisconsortium.com/pn-annotation/, release 2024-04-15), with filtering applied to include only E3 ubiquitin/UBL ligases, proteasome-associated proteins, and chaperones.

### Statistical analyses

Statistical analyses were performed using GraphPad Prism (v.10.1.0) for qRT-PCR results in Fig.S1A, using a two-tailed unpaired Welch’s t-test. For transcriptomics data, R (v4.4.1) was used with the glmQLFTest function in the edgeR (v4.2.2) package. For proteomics data, Perseus (v1.6.10.0) was used with two-tailed unpaired Student’s t-tests or Welch’s t-tests with a permutation-based false-discovery rate (FDR) of 0.05 and s0 = 0 (default). Gene Ontology (GO) enrichment analysis was performed using clusterProfiler (v4.12.6) in R, applying the Benjamini-Hochberg method (adjusted p < 0.05, q-value < 0.02). For more details on the statistical analyses and methods used in transcriptomics, proteomics, and Gene Ontology enrichment analysis, please refer to the corresponding sections above.

## Supporting information

Supplementary Data: Raw Data for Graphs and Omics Analyses

## Acknowledgements

This work was funded by Deutsche Forschungsgemeinschaft (DFG, German Research Foundation) SFB1430 - Project-ID 424228829 (to A.P., D.H., F.K., M.E. M.K.). J.S. is supported by the German Cancer Consortium (DKTK) and by the German Federal Ministry of Education and Research (BMBF; 01KD2206A/SATURN3). We thank Jenny Bormann and Dr. Sabrina Ninck from the Analytical Core Facility Essen (University of Duisburg-Essen) for their technical support in sample preparation for proteomics.

## Author contributions

H.T. designed and executed experiments, analyzed data and drafted the manuscript; F.K. and M.K. performed mass spectrometry; S.S.L. performed RNA sequencing; A.P., D.H., and J.S. edited the manuscript; M.E. outlined the work and wrote the manuscript with contributions from all authors.

## Data availability

The mass spectrometry proteomics data generated in this study have been deposited in the ProteomeXchange Consortium via the PRoteomics IDEntifications (PRIDE) Database repository with the dataset identifier PXD060306. During the review process the data can be accessed via a reviewer account (Password: 1uGdFCV4ICBJ). The RNA-seq data have been deposited in NCBI’s Gene Expression Omnibus database under accession codes GSE306539.

---

**Fig. S1.**
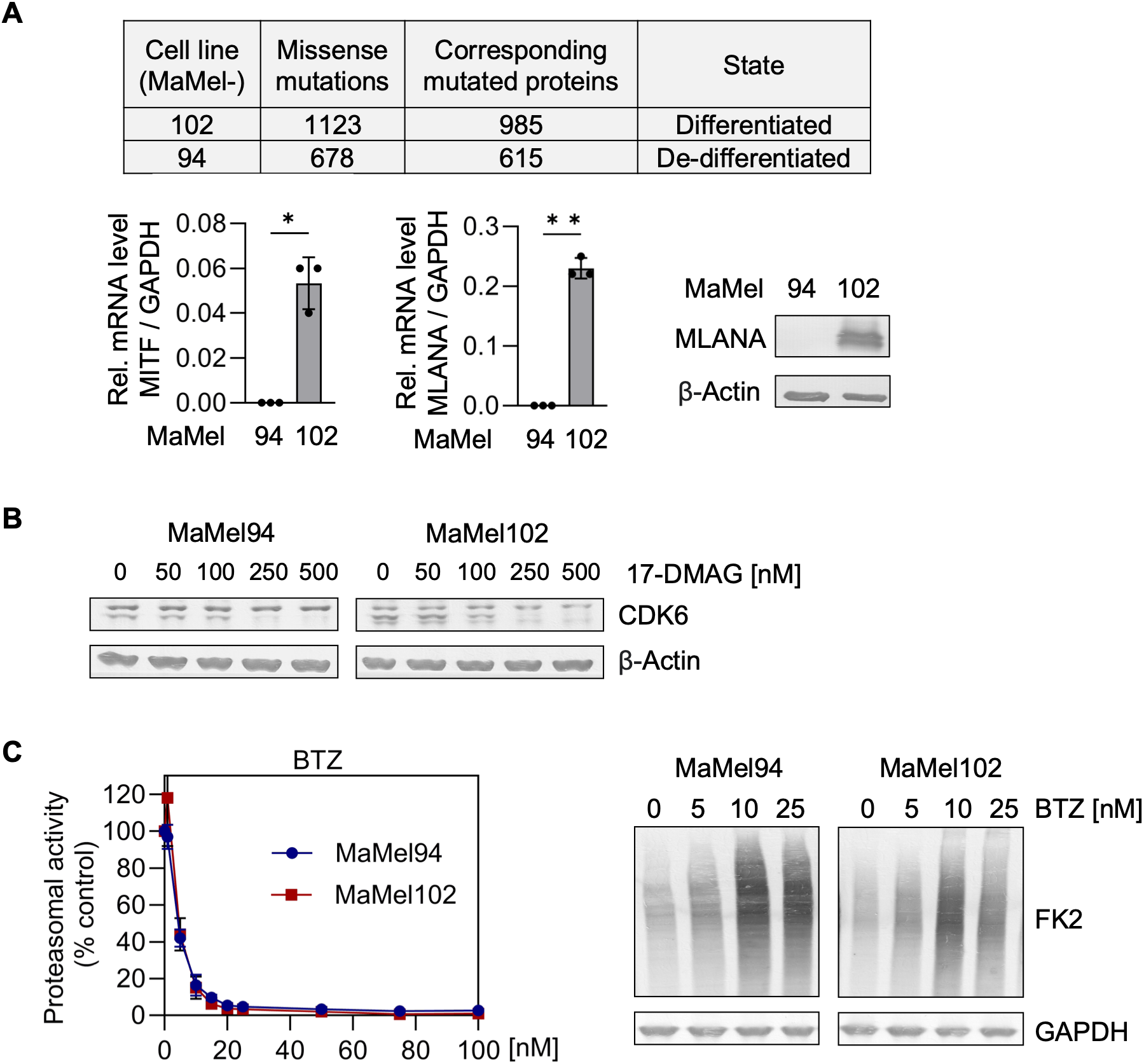
Related to Fig. 1. Assessment of melanoma differentiation state and efficacy of inhibitors. **(A)** Table summarizes the number of missense mutations and corresponding proteins in MaMel94 and MaMel102 cells. qRT-PCR analysis of melanoma differentiation markers MITF and MLANA was performed to assess their relative expression normalized to GAPDH. Data represent the mean ± SD values from three biological replicates. Statistical significance was determined using a two-tailed Welch’s t-test (*p ≤ 0.05, **p ≤ 0.01). Western blot analysis of MLANA supports the differentiation melanoma state; β-Actin was used as a loading control. **(B)** The efficacy of 17-DMAG as an HSP90 inhibitor was confirmed by Western blot analysis of CDK6, an HSP90 client protein. MaMel94 and MaMel102 cells were treated with 17-DMAG for 24 h at the indicated concentrations, with DMSO as a vehicle control. β-actin served as a loading control. **(C)** The efficacy of BTZ as a proteasome inhibitor was confirmed by a Proteasome-Glo assay (8 h treatment, left panel) and Western blot analysis of ubiquitinated proteins using the FK2 antibody (24 h treatment, right panel). GAPDH was used as a loading control for the Western blot.

**Fig. S2.**
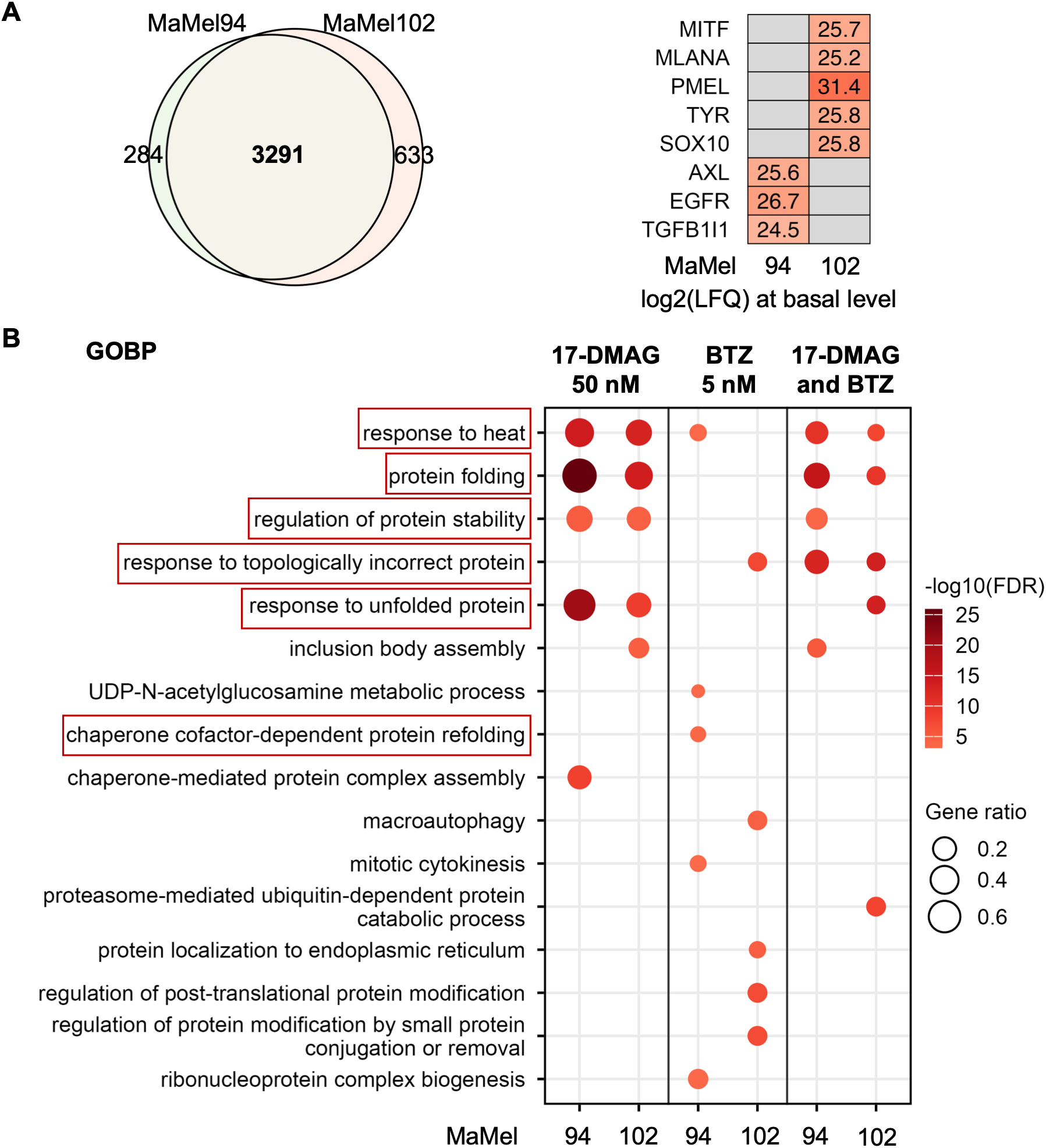
Related to Fig. 2. Proteomic analysis of melanoma cells treated with HSP90 and proteasome inhibitors. **(A)** Venn diagram shows the number of proteins identified in MaMel94 and MaMel102 in the proteomics analysis. Heatmap displays average log2(LFQ) values at basal level of melanoma differentiation markers (MITF, MLANA, PMEL, TYR, SOX10) and de-differentiation markers (AXL, EGFR, TGFB1I1). Gray indicates protein not detected. **(B)** Gene Ontology Biological Process (GOBP) enrichment analysis of differentially abundant proteins in each treatment condition shows strong enrichment of protein-folding–related processes across treatments and cell lines. Circle color intensity reflects significance, and size represents gene ratio. The top 5 enriched processes per treatment are shown.

**Fig. S3.**
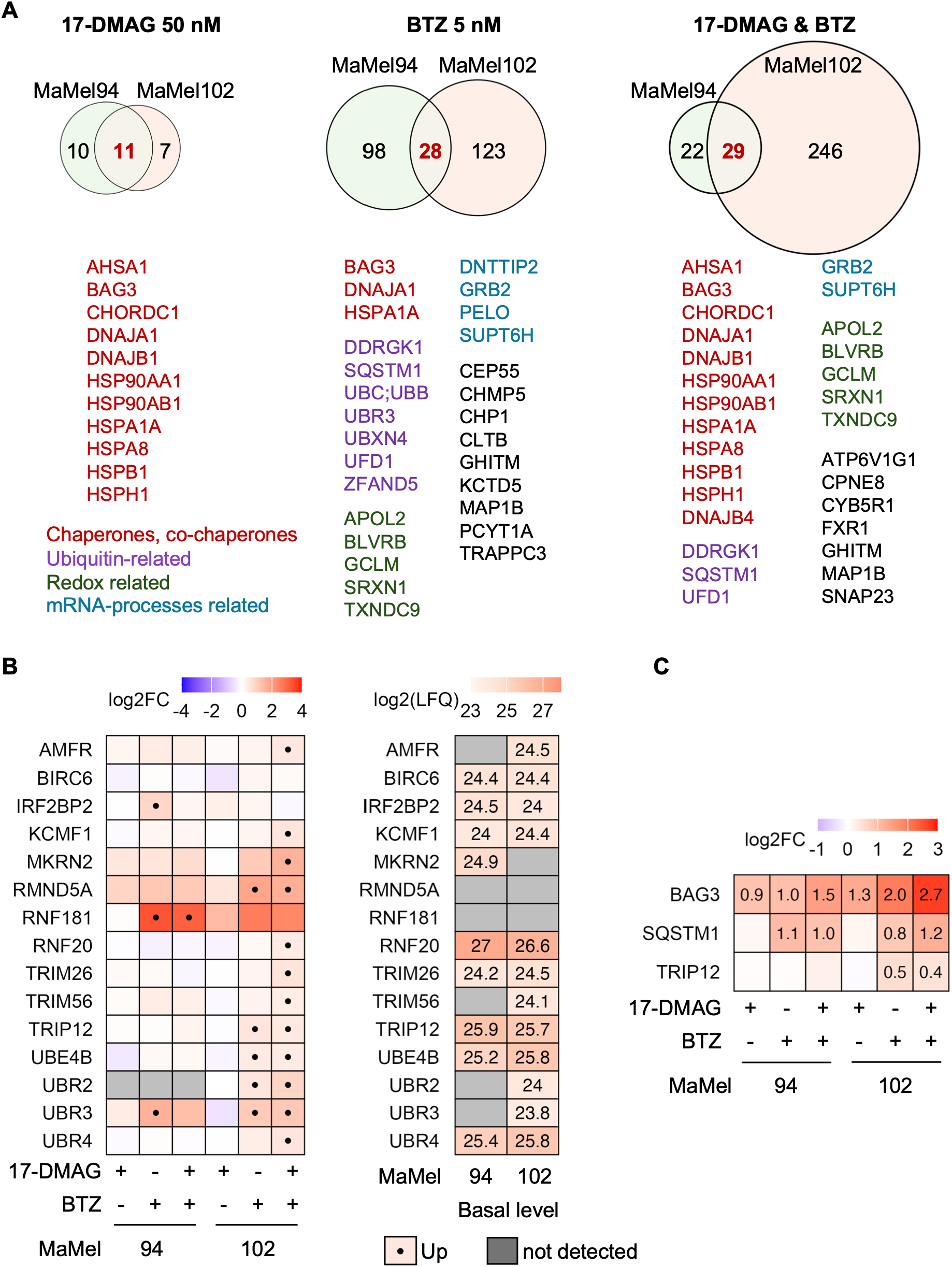
Related to Fig. 2. Proteomic analysis of melanoma cells treated with HSP90 and proteasome inhibitors. **(A)** Venn diagrams show the overlap of upregulated proteins between MaMel94 and MaMel102 across treatments, as identified by unpaired two-tailed t-tests with log2FC ≥ log2(1.3) and adjusted p-value < 0.05. Commonly upregulated proteins are listed for each treatment condition. Proteins sharing similar functions are highlighted by colors: red for “Chaperones, co-chaperones”, purple for “Ubiquitin-related”, green for “Redox related”, and blue for “mRNA-processes related”. **(B)** Heatmap of E3 ligases which were upregulated in at least one treatment condition in at least one cell line, showing log2FC compared to DMSO control from unpaired two-tailed t-tests (left panel) and average log2(LFQ) values at basal level (right panel). Gray indicates protein not detected by proteomics. Dot indicates E3 ligase that was upregulated (log2FC ≥ log2(1.3) and adjusted p-value < 0.05). **(C)** Heatmap of log2FC of proteins used for Western blot validation, with numerical values shown only for upregulated proteins (log2FC ≥ log2(1.3) and adjusted p-value < 0.05).

**Fig. S4.**
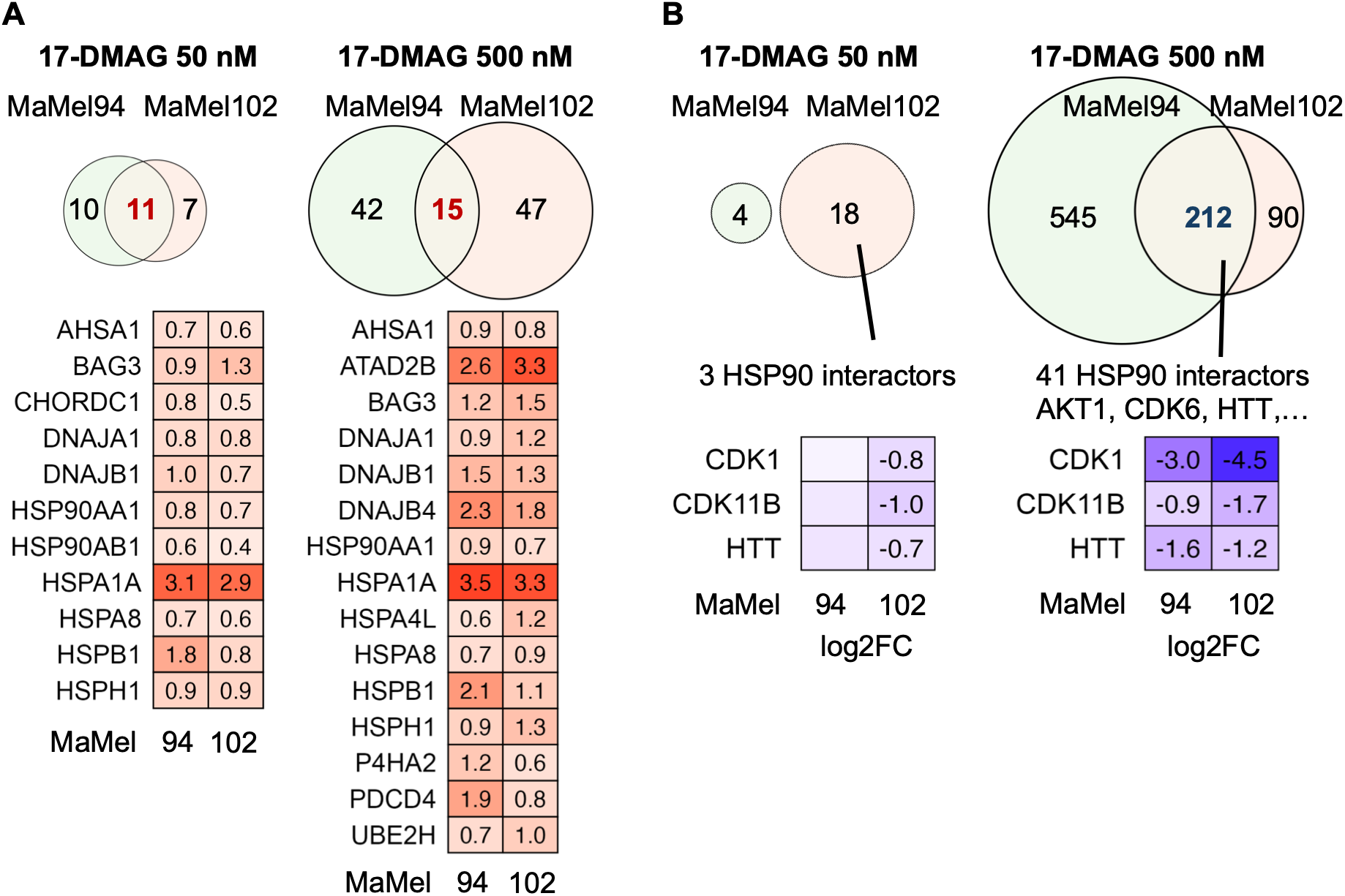
Related to Fig. 3. Differential protein abundance upon HSP90 inhibition with 17-DMAG at 50 nM vs 500 nM. **(A)** Venn diagrams and heatmaps show upregulated proteins shared between MaMel94 and MaMel102 upon 17-DMAG treatment at 50 nM and 500 nM. Proteins were considered upregulated if log2FC ≥ log2(1.3) and adjusted p-value < 0.05 for 50 nM treatment, or log2FC ≥ log2(1.5) and adjusted p-value < 0.05 for 500 nM treatment. Heatmap displays log2FC values. **(B)** Venn diagrams and heatmaps show downregulated proteins shared between MaMel94 and MaMel102 upon 17-DMAG treatment at 50 nM and 500 nM. Proteins were considered downregulated if log2FC ≤-log2(1.3) and adjusted p-value < 0.05 for 50 nM treatment, or log2FC ≤-log2(1.5) and adjusted p-value < 0.05 for 500 nM treatment. Heatmap displays log2FC values for selected HSP90 interactors, with numerical values shown only for downregulated proteins (log2FC ≤-log2(1.3) and adjusted p-value < 0.05).

**Fig. S5.**
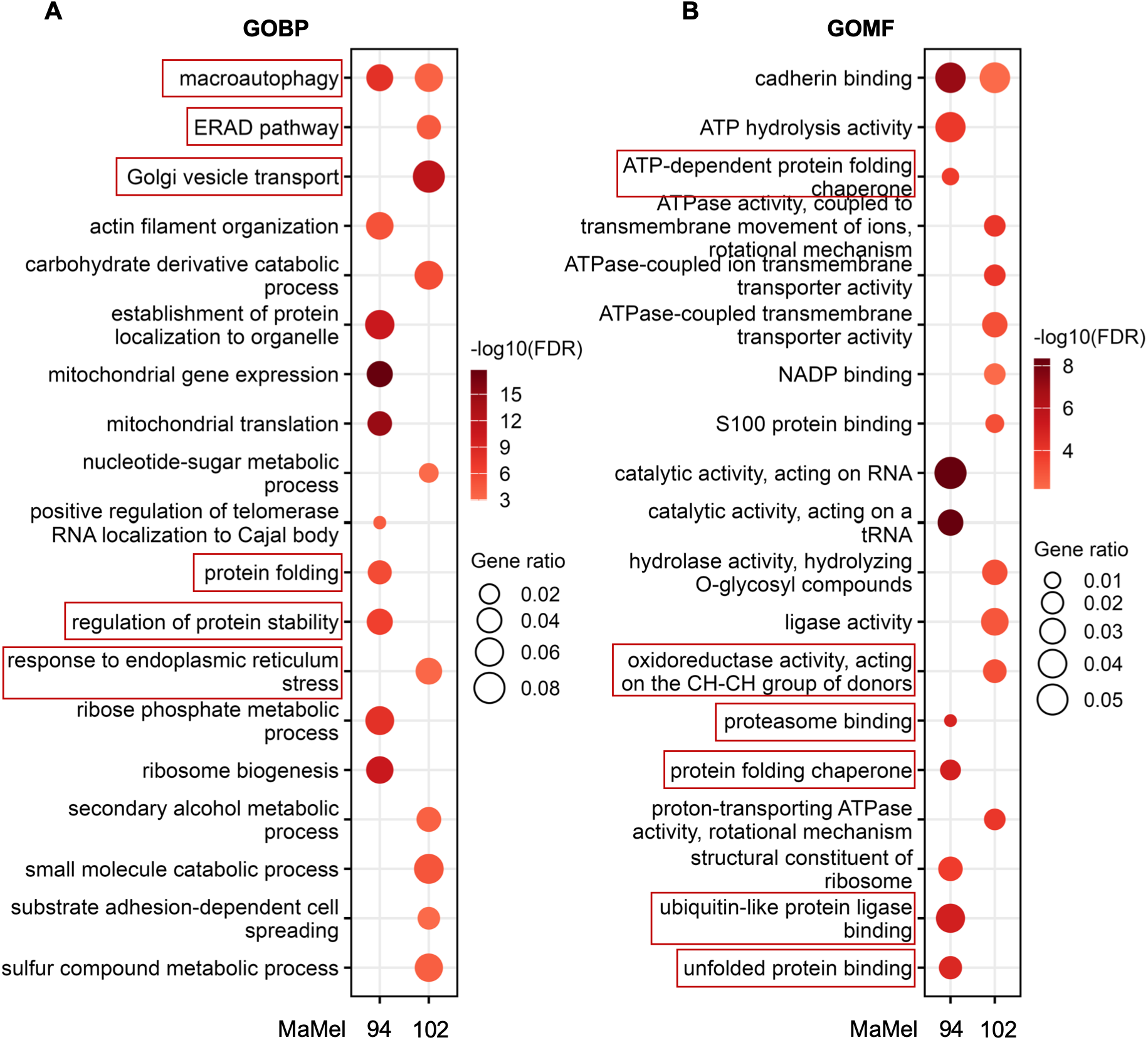
Related to Fig. 3. Top 10 enriched GOBP and GOMF terms among proteins downregulated following TRIP12 depletion in each cell line. Gene Ontology (GO) enrichment analysis was performed on proteins downregulated in each cell line following TRIP12 depletion (log2FC ≤-log2(1.5) and adjusted p-value < 0.05). The top 10 enriched Biological Processes (GOBP) and Molecular Functions (GOMF) per cell line are shown in **(A)** and **(B)**, respectively. Circle color intensity represents the significance of the enrichment, with darker colors indicating higher significance. Circle size is proportional to the gene ratio, reflecting the proportion of genes involved in each term.

**Fig. S6.**
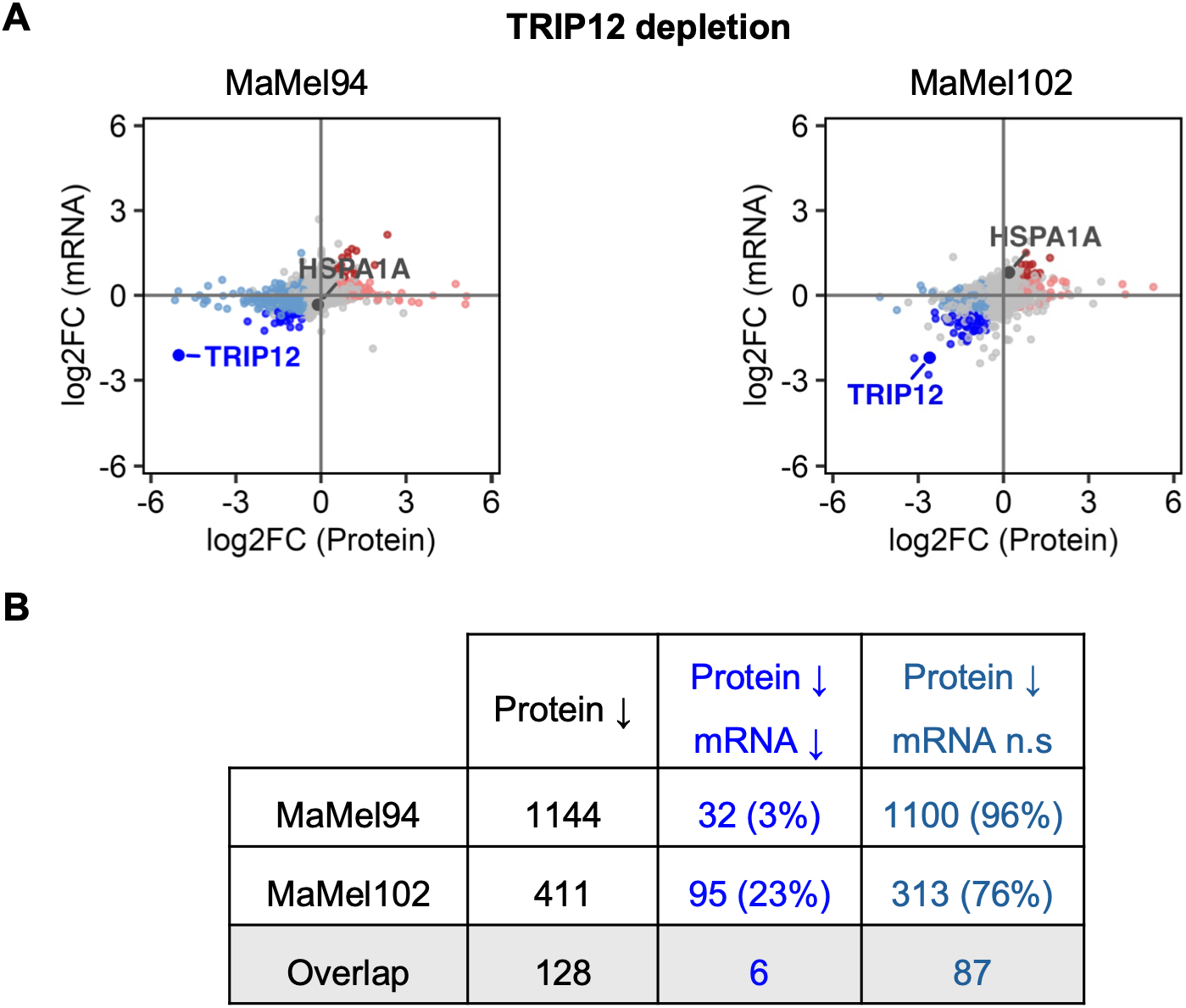
Related to Fig. 3. Comparison of protein and mRNA changes following TRIP12 depletion. **(A)** Scatter plots illustrate the correlation between protein and mRNA changes upon TRIP12 depletion in MaMel94 and MaMel102. The x-axis represents the log2 fold change (log2FC) of proteins, and the y-axis represents the log2FC of the corresponding mRNAs. The color scheme indicates the direction of change: red for upregulated proteins with upregulated mRNAs, pink for upregulated proteins with unchanged mRNAs, dark blue for downregulated proteins with downregulated mRNAs, and light blue for downregulated proteins with unchanged mRNAs. **(B)** Table summarizes the total number of downregulated proteins, as well as the number and percentage of protein–mRNA pairs with concordant changes in response to TRIP12 depletion. Protein–mRNA pairs are categorized as follows: dark blue for proteins downregulated with concurrent downregulation of their corresponding mRNAs, light blue for proteins downregulated with unchanged mRNA levels.

**Fig. S7.**
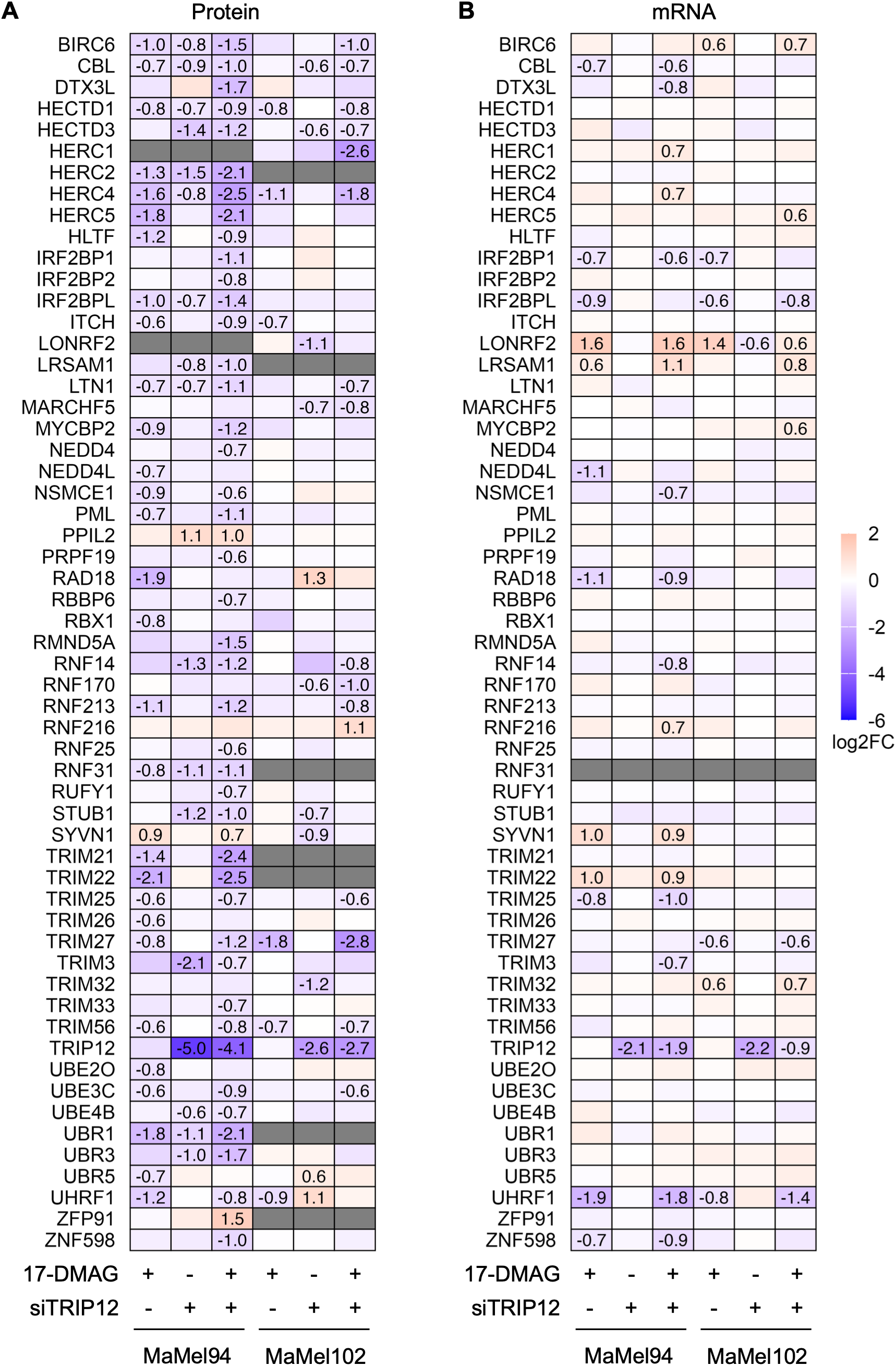
Related to Fig. 3D and Fig. 4A. E3 ligase protein and mRNA changes following TRIP12 depletion alone or combined with HSP90 inhibition. Heatmap of log2FC values for E3 ligase proteins **(A)** and their corresponding mRNAs **(B)** in MaMel94 and MaMel102 in response to HSP90 inhibition alone, TRIP12 depletion alone, or combined HSP90 inhibition and TRIP12 depletion. Log2FC values are shown only for proteins and genes that were differentially abundant under the corresponding condition (absolute log2FC ≥ log2(1.5), adjusted p-value < 0.05). Gray indicates protein or gene not detected in the respective cell line.

**Fig. S8.**
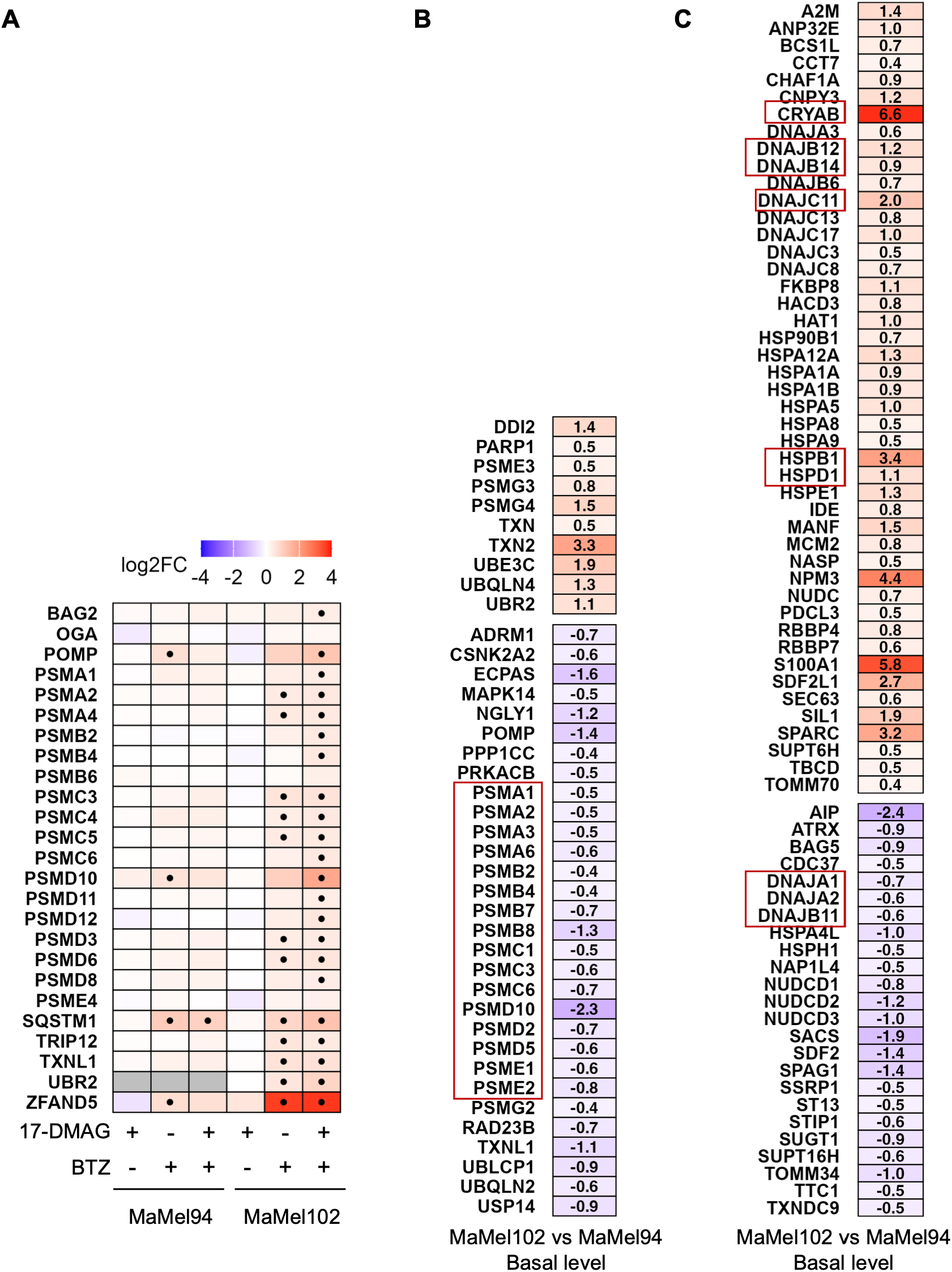
Related to Fig. 2B. Basal proteostasis-factor abundance is associated with differential responses to proteasome inhibition in MaMel94 and MaMel102. **(A)** Heatmap of log2FC values for proteasome-associated proteins upregulated in response to at least one treatment condition (17-DMAG 50 nM and/or BTZ 5 nM) in at least one cell line. Gray indicates protein not detected by proteomics. Dot indicates proteins upregulated under the corresponding condition (adjusted p-value < 0.05 and fold change ≥ 1.3). **(B, C)** Heatmaps of log2FC values for proteasome-associated proteins (B) and chaperone-related proteins (C) at basal levels, showing MaMel102 relative to MaMel94.

**Table S1.** Primary antibodies.

| Primary antibody | Host organism | Dilution (WB) | Source | Identifier |
|---|---|---|---|---|
| Actin (EPR16770) | rabbit monoclonal | 1:20000 | Abcam | Cat# ab198991 |
| BAG3 | rabbit polyclonal | 1:10000 | Proteintech | Cat# 10599-1-AP |
| CDK6 | rabbit polyclonal | 1:1500 | Proteintech | Cat# 14052-1-AP |
| FANCD2 | rabbit polyclonal | 1:2500 | Proteintech | Cat# 28619-1-AP |
| FANCI | mouse monoclonal | 1:10000 | Proteintech | Cat# 67304-1-Ig |
| GAPDH (14C10) | rabbit monoclonal | 1:10000 | Cell Signaling Technology | Cat# 2118 |
| HSP90AA1 | rabbit polyclonal | 1:20000 | Proteintech | Cat# 13171-1-AP |
| HSPA1A | rabbit polyclonal | 1:25000 | Proteintech | Cat# 10995-1-AP |
| HSPB1/ HSP27 | rabbit polyclonal | 1:25000 | Proteintech | Cat# 18284-1-AP |
| KDM2A | rabbit polyclonal | 1:2000 | Proteintech | Cat# 24311-1-AP |
| MLANA/MART-1 | mouse monoclonal | 1:5000 | Zytomed Systems | Cat# MSG056 |
| Multi-Ubiquitin (FK2) | mouse monoclonal | 1:1000 | MBL | Cat# D058-3 |
| MX1 | rabbit monoclonal | 1:20000 | Proteintech | Cat# 82677-1-RR |
| OGFR | rabbit monoclonal | 1:20000 | Proteintech | Cat# 85998-1-RR |
| p62/SQSTM1 | rabbit polyclonal | 1:10000 | Proteintech | Cat# 18420-1-AP |
| TRIP12 | rabbit polyclonal | 1:1000 | Proteintech | Cat# 25303-1-AP |

**Table S2:**
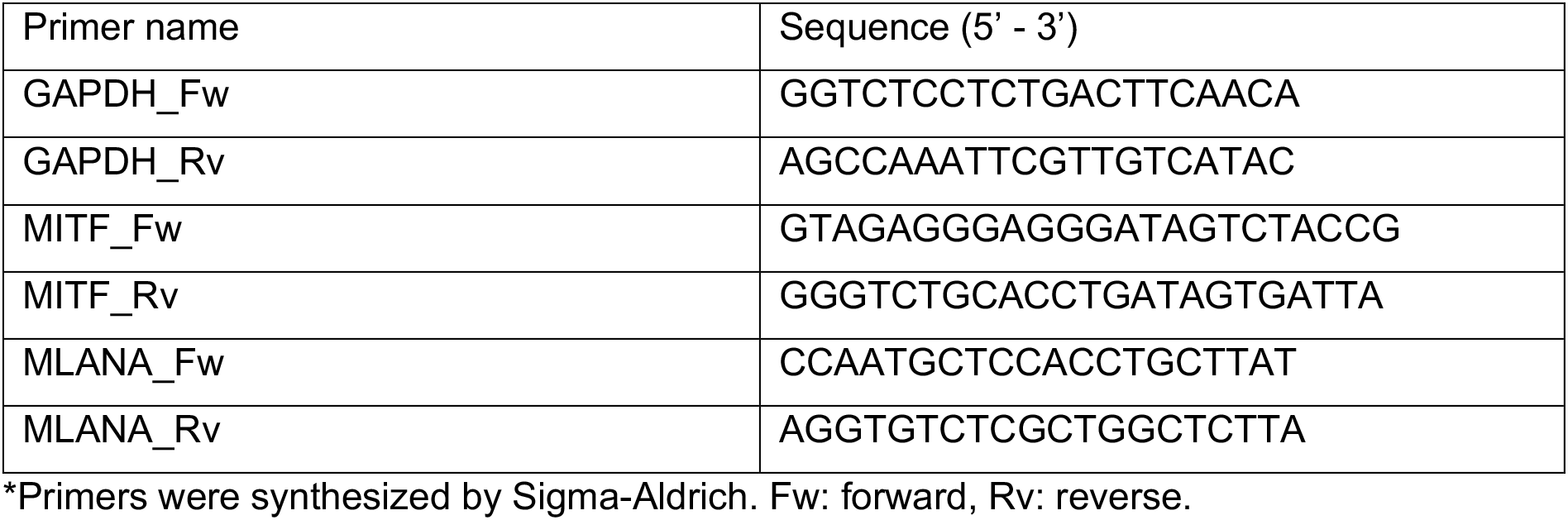
Primer sequences for qRT-PCR.

